# Reconstructed signaling and regulatory networks identify potential drugs for SARS-CoV-2 infection

**DOI:** 10.1101/2020.06.01.127589

**Authors:** Jun Ding, Jose Lugo-Martinez, Ye Yuan, Jessie Huang, Adam J. Hume, Ellen L. Suder, Elke Mühlberger, Darrell N. Kotton, Ziv Bar-Joseph

## Abstract

Several molecular datasets have been recently compiled to characterize the activity of SARS-CoV-2 within human cells. Here we extend computational methods to integrate several different types of sequence, functional and interaction data to reconstruct networks and pathways activated by the virus in host cells. We identify key proteins in these networks and further intersect them with genes differentially expressed at conditions that are known to impact viral activity. Several of the top ranked genes do not directly interact with virus proteins. We experimentally tested treatments for a number of the predicted targets. We show that blocking one of the predicted indirect targets significantly reduces viral loads in stem cell-derived alveolar epithelial type II cells (iAT2s).

**Software and interactive visualization:** https://github.com/phoenixding/sdremsc

## Introduction

To fully understand the impact of the novel coronavirus, SARS-CoV-2, which causes the COVID-19 pandemic, requires the integration of several different types of molecular and cellular data. SARS-CoV-2 is known to primarily impact cells via two viral entry factors, ACE2 and TMPRSS2 [1]. However, much less is currently known about the virus activity once it enters lung cells. Similar to other viruses, once it enters the cell response to SARS-CoV-2 leads to the activation and repression of several pro- and anti-inflammatory pathways and networks [2]. Predictive mechanistic understanding about the activity of these pathways, the key proteins that participate in them and the regulatory networks that these pathways activate is the first step towards identifying useful drug targets.

Recently, several studies provided information about various aspects of the molecular activity of SARS-CoV-2. These include studies focused on inferring virus-host interactions [3, 4], studies focused on the viral sequence and mutations [5], studies profiling expression changes following infection [6] and studies identifying underlying health conditions that lead to an increased likelihood of infection and death [7, 8]. While these studies are very informative, each only provides a specific viewpoint regarding virus activity. By integrating all of this data in a single interaction network we can enhance each of the specific data types and reconstruct the networks utilized by the virus and host following infection.

Several unsupervised methods have been developed for inferring molecular networks from interaction and expression data [9, 10]. However, these usually cannot link the expression changes observed following infection to the viral proteins that initiate the responses. We have previously developed SDREM [11, 12] to link sources (viral proteins and their human interactors) with targets (genes that are activated / repressed following infection) in order to reconstruct regulatory and signaling networks and identify key proteins that mediate infection signals. SDREM combines Input-Output Hidden Markov models (IOHMM) with graph directionality assignments to link sources and targets. It then performs combinatorial network analysis to identify key nodes that can block the signals between sources and targets and ranks them to identify potential targets for treatments. Here we extended SDREM in a number of ways to reconstruct SARS-CoV-2 pathways. First, we developed methods that allow SDREM to use time series single cell RNA-Seq (scRNA-Seq) data. We also extended SDREM so that (static) phosphorylation data can be used to refine the selected pathways. Finally, we improved the scoring function by intersecting top ranked SDREM genes with genes identified by complementary gene expression and screen studies.

The extended method was applied to three gene expression datasets profiling responses to SARS-CoV-2 in bulk and single cells. As we show, the networks reconstructed using these datasets were in very good agreement and also agree with independent proteomics studies. The networks identified several relevant genes as potential targets. Intersecting the list of top-scoring SDREM genes with genes identified as differentially expressed (DE) in several of conditions associated with COVID-19 severity further narrows the list of candidate targets. Many of the top predictions are not directly interacting with viral proteins and so cannot be identified without integrating several different data types. We tested a number of the predictions and demonstrated the effectiveness of one of the predicted drugs, bortezomib, at inhibiting SARS-CoV-2 infection in human induced pluripotent stem cell-derived alveolar epithelial type II cells (iAT2s).

## Results

We integrated several condition specific and general molecular datasets to reconstruct infection pathways for SARS-CoV-2 in lung cells. To reconstruct activated and repressed pathways and their potential impact on viral load we start with host proteins known to interact with virus proteins [3, 4], and attempt to link them through signaling pathways to dynamic regulatory networks reconstructed using gene expression profiling following viral infection [13, 14, 15]. Next, our method ranks all proteins based on their participation in pathways that link sources (proteins directly interacting with the virus) and targets (expressed genes) and selects top ranking pathways. These are used to reconstruct networks which contain proteins and interactions that are likely to play key roles in either increasing or decreasing viral loads. We further intersect top ranked proteins with a large cohort of lung expression data from conditions that are known to impact SARS-CoV-2 infection and severity and with functional SARS-CoV-2 screens of host genes.

### Reconstructed signaling and regulatory network from bulk transcriptomics

We used our revised SDREM method to reconstruct signaling and regulatory networks activated following SARS-CoV-2 infection. We first combined virus-host and host-host protein interaction data with bulk time series expression data of lung epithelial cells infected by SARS-CoV-2 (Methods). The reconstructed network is presented in Supplementary Figure S1. The signaling part contained 252 proteins, of which 203 are source proteins (host proteins that interact with viral proteins) and 49 are proteins that do not directly interact with viral proteins. These 49 proteins included 34 TFs that directly regulate the expression downstream target genes and 15 internal (signaling) proteins which serve as the intermediate between the source proteins and the TFs. We first examined the 203 source proteins selected by the method. These represent only 17.7% of the 1148 human proteins that were experimentally identified as interacting with virus proteins (and that served as input to the method). The most enriched GO term (using ToppGene [16]) for the 203 selected source proteins is ‘viral process’ (FDR=2.178e-14). In contrast, no GO category related to viruses is found to be significant for the full set of 1148 source proteins. This indicates that by integrating several diverse datasets, our extended SDREM model is able to zoom in on the most relevant source proteins from the two studies.

The 15 internal proteins are also enriched for relevant functions including ‘cellular response to steroid hormone stimulus (FDR=1.153e-9)’ which is also significantly associated with the 34 target TFs (FDR=1.666e-12). The combined set of 49 non-source proteins are also significantly enriched with sex hormone response related functions. The role of sex hormone in COVID-19 was recently reported [17].

We further validated the proteins identified by SDREM by comparing them to 543 proteins that were determined to be phosphorylated following infection with SARS-CoV-2 [18]. We observed a significant overlap between SDREM selected proteins and the list of phosphorylated proteins (19 out of 252 proteins are phosphorylated: hyper-geometric test p-value=6.35e-5).

### Integrating phosphorylation data to reconstruct networks

While the phosphorylation data can serve as a validation, we can also use it as an input to increase the prior for including a protein in the reconstructed network (Methods). Given the significant intersection between the networks reconstructed without such data and the phosphorylation data we next reconstructed networks that use this data in addition to the protein-protein, protein-DNA interaction and time series expression data.

Networks learned with this data included a total of 261 proteins with 204 source proteins 17 internal proteins and 40 TFs (Figure 1). We again observe ‘viral process’ as the most enriched category for the source proteins (Supplementary Table S1). (FDR=1.173e-10). The 17 internal proteins are significantly enriched with transcription relevant GO terms such as “negative regulation of transcription, DNA-templated (FDR=2.772e-7)” and sex hormone stimulus relevant functions such as “intracellular steroid hormone receptor signaling pathway (FDR=2.772e-7)”. The 40 target TFs are enriched with “positive regulation of transcription by RNA polymerase II (FDR=1.066E-39)” and “response to hormone (FDR=2.738E-12)”, which are consistent with the function of internal proteins.

**Figure 1:**
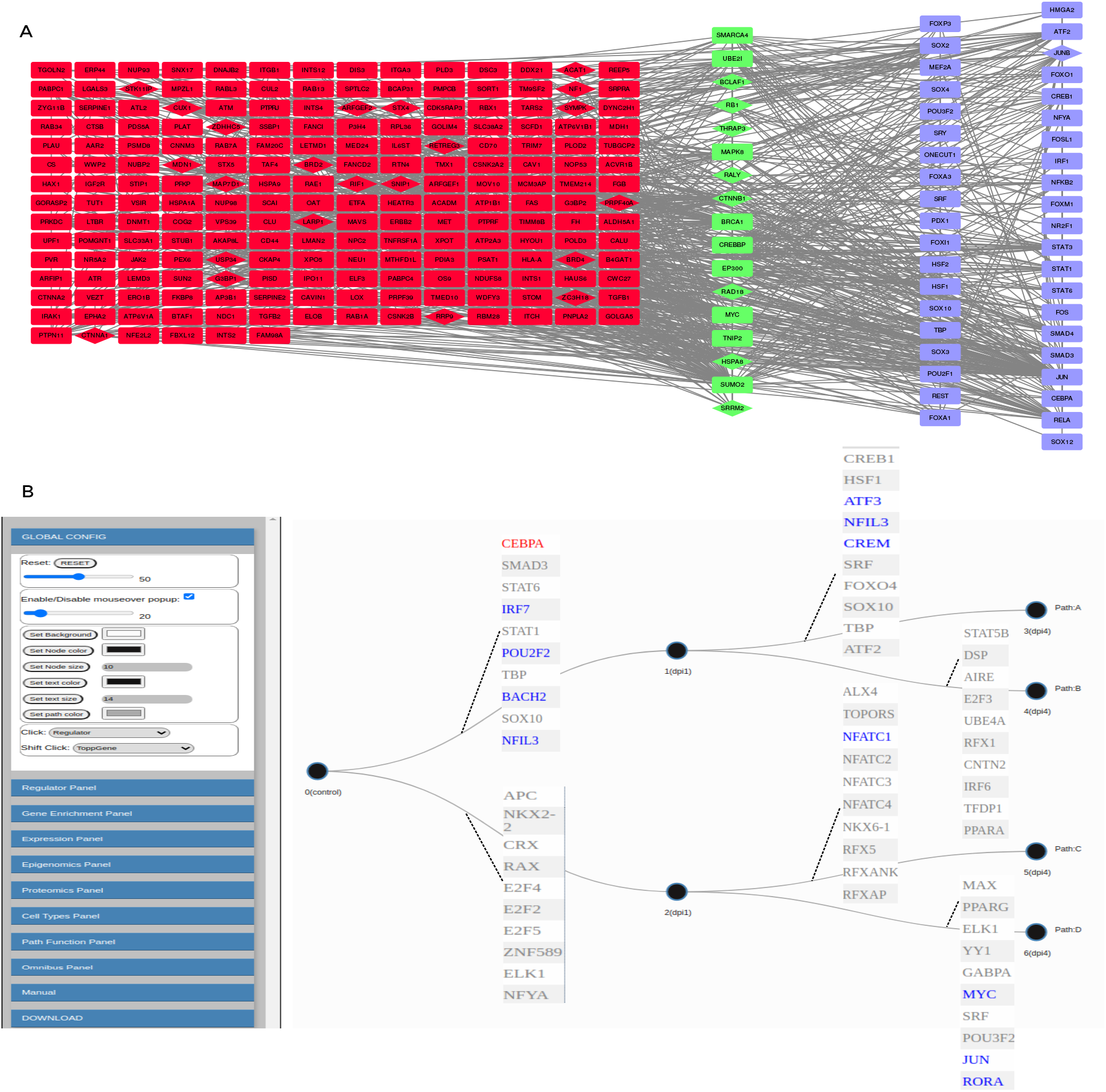
SDREM predicted signaling network(A) and regulatory network (B) for the time-series RNA-seq dataset (with using protein phosphorylation data). (A) **signaling network reconstructed from the time-series single-cell RNA-seq data**. Red nodes denote sources, green nodes are signaling proteins and blue nodes are TFs associated with regulating the DREM portion of the model. The overlap between the networks is discussed in Results. (B) **The regulatory part of the network** Each path represents a group of genes that share similar expression profiles. The table presented next to each edge indicates the set of transcription factors (TFs) that are predicted to regulate the expression of genes assigned to this path. Red font indicates TF expression is significantly down-regulated, while blue represents up-regulated TFs. TF with stable expression (inferred to be post-transcriptionally or post-translationally regulated) are marked as Gray.

In addition to identifying proteins that play a major role in key pathways, SDREM can also be used to identify pairs of proteins that, together, control a significant number of pathways (i.e., pairs that are expected to have the largest impact in a double KO experiment, Methods). For the learned network SDREM also identified 28712 protein pairs. These protein pairs are composed of 243 distinct proteins (203 sources, 17 internal, 22 TFs), which slightly differ from the proteins identified based on their individual rankings (243/261=93.% single knock-out proteins are also found in double knock-out analysis). See Supplementary Table S1 for the complete list of protein pairs.

### Reconstructed signaling and regulatory network from single-cell transcriptomics

To further narrow down the list of key signaling and regulatory factors involved in SARS-CoV-2 response we next used SDREM to analyze time series Calu-3 scRNA-Seq SARS-CoV-2 infection data [14](Methods). The signaling part reconstructed for this data contained 244 proteins, of which 172 are source proteins (host proteins that interact with viral proteins) and 72 are proteins that do not directly interact with viral proteins. These 72 proteins included 49 TFs that directly regulate the expression downstream target genes and 23 internal (signaling) proteins which serve as the intermediate between the source proteins and the TFs. We found that the majority of the 244 proteins (159/65.2%, p-value=2.26e-12) are shared between the single-cell and bulk reconstructed networks (135, source proteins 14 TFs, and 10 internal proteins). Figure 2 presents the conserved signaling network at the intersection of both networks. The most enriched GO term (using ToppGene) for the 135 selected source proteins is ‘viral process’ (FDR=1.01e-11). The enriched GO terms for the 24 non-source proteins include ‘regulation of transcription’(FDR=3.23e-19), and ‘response to cytokine’ (FDR=6.250E-15). Please refer to Supplementary Table S2 for the detailed GO terms.

**Figure 2:**
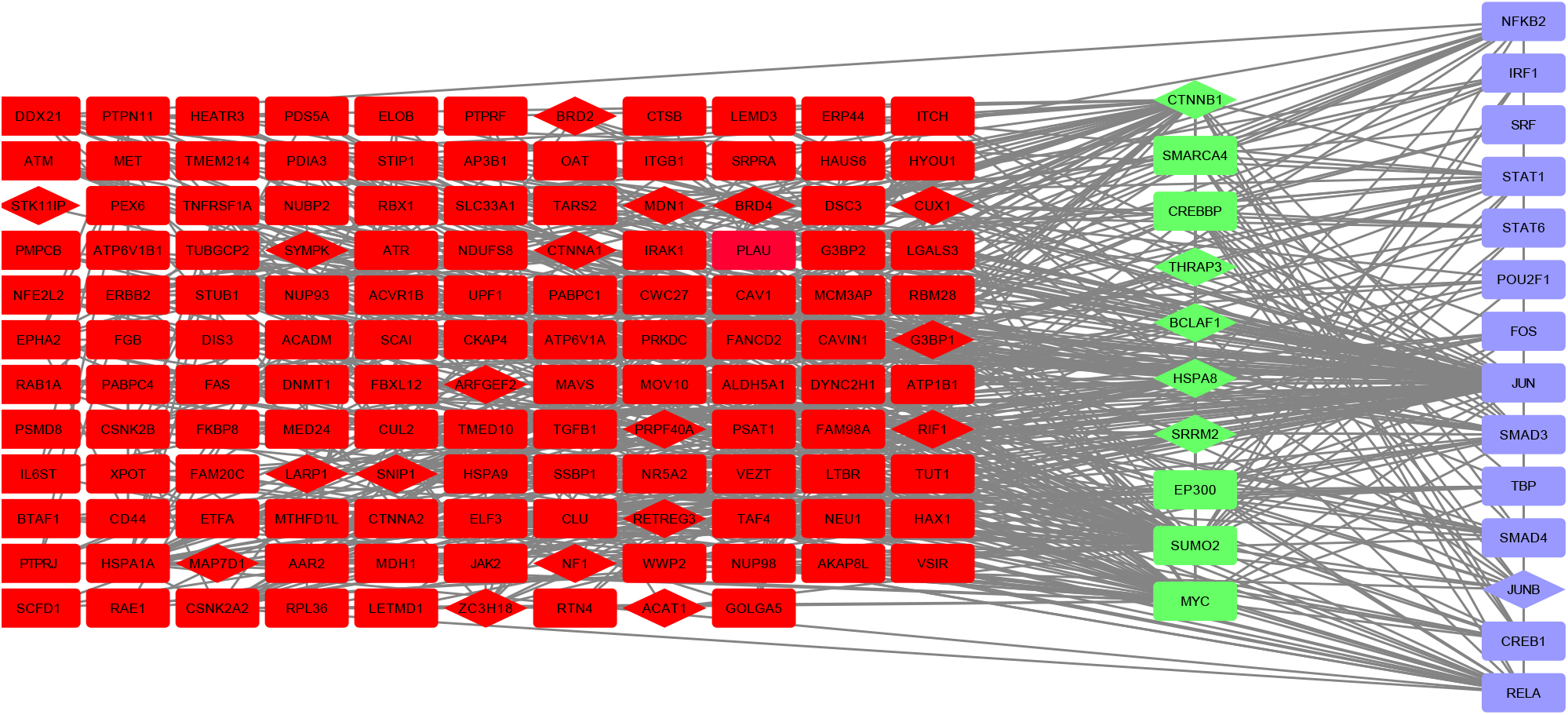
The conserved signaling network between bulk and single-cell SDREM reconstructed networks. Notations in this figure are the same as in Figure 1A.

We also used SDREM to analyze another time-series scRNA-Seq infection data using human airway epithelium cells [15](Methods). For this data the reconstructed network is composed of 288 proteins, of which 225 are source proteins, 15 are internal proteins, and 48 are transcription factors. Again, the reconstructed network was in good agreement with the bulk data network discussed above (74.8% overlap in identified proteins). See Supplementary Information for complete list of proteins identified for this data and for their enrichment GO terms.

### Intersection with disease genes, RNAi or CRISPR knockdown studies

We studied the activity of SDREM identified genes in a set underlying health conditions that were determined to impact SARS-CoV-2 infection and mortality rates, including hypertension [19], age [20] and gender [20], See Supplementary Table S3 for the complete set of conditions we considered. The intersection results are presented in Supplementary Figures S3-S10. We identified several genes in the overlap of the set of conditions we tested and the SDREM set (p value = 2.63e-6, hypergeometric distribution). Some of the genes in the intersection are TFs that are expressed in many tissue (e.g., JUN and FOS) and are also known to have important roles in lung cells. CAV 1 is another interesting gene which is essential during the acute lung injury [21]. More generally the list of genes is enriched for GO terms that include ‘cellular response to chemical stimulus’ (p value = 1.80e-11), ‘viral process’ (p value = 2.37e-10), transcription and metabolism relevant functions. We next compared the 261 proteins identified by SDREM from bulk RNA-seq expression with RNAi and CRISPR screens for multiple coronaviruses (Methods). We identified 18 proteins (p value = 2.49e-4) which have been previously shown to affect coronavirus load in RNAi screen experiments (Table 1). It is worth noting that 5 genes on this list (ATM, ATP6V1A, CAV1, SMAD3 and UBE2I) are also identified as condition related genes (p value = 8.93e-3) and thus appear in all relevant datasets we analyzed (network analysis, condition related and RNAi). The smaller list of top 1000 protein pairs identified by SDREM (168 proteins) includes 10 proteins (p value = 1.52e-2) which have been annotated to alter coronavirus replication across different RNAi experiments. Of these 10 proteins, RAB7A is a potentially interesting target [22, 23, 24]. RAB7A is a lysosomal-endosomal protein that is found in alveolar epithelial type 2 (AT2) cells. RAB7A has an important role in disease pathogenesis and is part of both the endosomal and lamlellar body-multivesicular body organelles whose normal function is required for proper surfactant packaging and secretion in the lung [25]. In addition to the 5 genes previously identified as condition-associated by the single ranking method the pairs method also identified EPHA2 as a condition related gene (p value = 3.16e-2). Table 1 summarizes the identified proteins supported by RNAi or CRISPR experiments. In a similar manner, we also analyzed the 170 proteins identified by SDREM in both of the scRNA-Seq reconstructed networks. Table 1 lists the 12 proteins identified from this set in RNAi or CRISPR screen experiments (p value = 2.12e-3). 11 of these proteins (∼92%) are identified by all three reconstructed networks (bulk, sc1 and sc2 in Table 1). Finally, the intersection between the top 1000 protein pairs in each reconstructed network identified by SDREM from single-cell data produced 285 pairs (composed of a total of 88 unique proteins). This list includes 6 proteins (∼50%) identified from the set of RNAi and CRISPR proteins (p value = 3.06e-2).

**Table 1:** Summary of proteins supported by RNAi or CRISPR screen experiments identified by SDREM from time-series bulk and single-cell SARS-CoV-2 expression data sets. For each protein, we list RNA-seq experimental evidence [‘Bu’ for bulk, ‘Sc1’ (Calu-3 cells) or ‘Sc2’ (human airway epithelium cells) for single-cell experiments, and ‘All’ for bulk and single-cell experiments], whether it is a known interactor of a SARS-CoV-2 protein and a brief description of the experimental impact on coronavirus load. Table enumerates each protein previously reported to affect coronavirus load in RNAi or CRISPR screen experiments for the set of proteins identified by SDREM from SARS-CoV-2 data (p values: 2.49e-4 for bulk data set; 9.28e-4 and 2.97e-4 for each single-cell data set, respectively). Proteins also listed in the top 1000 protein pairs identified by SDREM from either bulk or single-cell data sets are shown in boldface (p values: 1.52e-2 for bulk data; 8.97e-2 and 6.43e-3 for Sc1 and Sc2, respectively).

| Gene name | RNA-seq | SARS-CoV-2 interactor? | RNAi supported effect |
| --- | --- | --- | --- |
| POU3F2 | Bu | N | decreased IBV-CoV replication |
| **UBE2I** | Bu | N | decreased IBV-CoV replication |
| ATP6AP1 | Sc1 | Y | decreased SARS-CoV-2 replication |
| CAV2 | Sc1 | Y | decreased IBV-CoV replication |
| CSNK2A1 | Sc1 | N | decreased SARS-CoV replication |
| GBF1 | Sc2 | Y | decreased SARS-CoV-2 replication |
| **NFKB1** | Sc2 | N | decreased MERS-CoV replication |
| TTC27 | Sc2 | Y | decreased SARS-CoV replication |
| **SMARCA4** | Bu, Sc1 | N | conferred resistance to SARS-CoV-2 |
| NPC2 | Bu, Sc2 | Y | decreased SARS-CoV-2 replication |
| PISD | Bu, Sc2 | Y | decreased IBV-CoV replication |
| **RAB7A** | Bu, Sc2 | Y | affected MHV-CoV fusion<br>decreased SARS-CoV-2 load |
| **VPS39** | Bu, Sc2 | Y | affected MHV-CoV fusion |
| **ACVR1** | Sc1, Sc2 | Y | decreased SARS-CoV replication |
| **ACVR1B** | All | Y | decreased SARS-CoV replication |
| **ATM** | All | Y | decreased SARS-CoV-2 load |
| ATP6V1A | All | Y | decreased SARS-CoV-2 replication |
| **CAV1** | All | Y | decreased IBV-CoV replication |
| DYNC2H1 | All | Y | decreased MHV expression |
| **EPHA2** | All | Y | decreased SARS-CoV replication |
| G3BP2 | All | Y | decreased SARS-CoV-2 load |
| MDH1 | All | Y | decreased IBV replication |
| **RBX1** | All | Y | decreased IBV-CoV replication |
| **SMAD3** | All | N | conferred resistance to SARS-CoV-2 |
| **SMAD4** | All | N | conferred resistance to SARS-CoV-2 |

### Potential treatments for predicted genes

We looked for potential treatments for 107 total proteins (48 from bulk analysis as well as 67 and 53 from single-cell analysis on infected Calu-3 cells and human airway epithelium cells, respectively) identified at the intersection of top ranked SDREM and underlying condition genes. Supplementary Table S4 lists 34 human proteins that we identified as potential drug targets using curated databases of bioactive molecules such as ChEMBL, Pharos and ZINC. Supplementary Table S4 also provides the full list of chemical associations to human proteins identified as potential drug targets. Among these potential drug targets, 59% (i.e., 20 out of 34) are not characterized as SARS-CoV-2 interactors (i.e., non-source in our networks). Additionally, among the 34 human proteins, 16 proteins (> 47%) are identified separately from bulk data and at least one single-cell RNA-seq data set, including 7 proteins identified separately by all three data sets. It is worth highlighting that Gordon *et al*. [3] tested antiviral activity of 47 compounds targeting known SARS-CoV-2 interactors. Of these drug targets tested for inhibition of viral infection, only BRD2 and CSNK2A2 are also identified in our list. They reported viral inhibition for CSNK2A2 with compound Silmitasertib (CX-4945), whereas the antiviral activity results for BRD2/4 across 6 different compounds were inconclusive. In order to further analyze the SDREM reconstructed networks, we looked for approved drugs associated with the top 50 proteins identified in the intersection between bulk and single-cell SDREM reconstructed networks. Table 2 lists the resulting 5 proteins associated with at least one approved drugs, including the SDREM-based minimum rank between the bulk and single-cell lists and whether or not it is a known SARS-CoV-2. Interestingly, 40% (i.e., 2 out of 5) are not characterized as SARS-CoV-2 interactors which are ranked higher than known SARS-CoV-2 interactors.

**Table 2:** Summary of proteins associated with at least one approved drug and identified by SDREM from both time-series bulk and single-cell SARS-CoV-2. For each human protein, we show SDREM-based minimum rank between the bulk and single-cell lists, whether or not it is a known SARS-CoV-2 and a pair of associated approved drug names derived from ChEMBL25, IUPHAR/BPS Guide to Pharmacology, Pharos, or ZINC.

| Gene name | SDREM minRank | SARS-CoV-2 interactor? | Approved drug names |
| --- | --- | --- | --- |
| RELA | 11 | N | bortezomib |
| NFKB2 | 26 | N | bortezomib |
| DNMT1 | 31 | Y | azacitidine; decitabine |
| BRD4 | 32 | Y | alprazolam ; fedratinib |
| ERBB2 | 45 | Y | afatinib; neratinib |

### Experimental results

Following the identification of potential treatments based on SDREM analysis, we tested the ability of four FDA-approved drugs (Table 2) to inhibit SARS-CoV-2 infection the same cells that were used for the bulk expression data we analyzed (Figure 3). Multiple concentrations of each of these drugs were tested for their ability to inhibit SARS-CoV-2 infection of human induced pluripotent stem cell-derived alveolar type II cells (iAT2s) plated in air-liquid interface (ALI) cultures as described previously [13]. Three of the drugs showed mild to no efficacy in blocking infection with a recombinant SARS-CoV-2 clone expressing an mNeonGreen reporter replacing ORF7a (SARS-CoV-2-mNG) [26]. However, the fourth, bortezomib, appeared to reduce infection at both 1 and 10 *μ*M (Figure 3B). To test whether bortezomib treatment of iAT2s might also reduces infections with wild-type SARS-CoV-2, we repeated these tests and found statistically significant dose-response effects with moderate inhibition at 100 nM bortezomib and pronounced inhibition at higher concentrations as observed both by immunofluorescence and RT-qPCR quantitation of expression of SARS-CoV-2 nucleocapsid (N1) (Figure 3D-F). NFKBIA, a target gene of the canonical NF-*κ*B pathway whose mRNA expression levels correlate with pathway activity, was significantly down-regulated after treatment with 100 nM to 10 *μ*M of bortezomib, confirming the inhibitory effect of bortezomib on NF-*κ*B activity and suggesting a potential role in treating SARS-CoV-2 infections. Following these results we have further analyzed the RELA / NFKB molecular subnetwork that served as the basis for the high ranking assigned to this protein by SDREMs. Figure 4 presents genes in this subnetwork and their functional annotations. The centrality of RELA in our model is highlighted by the size of its sub-network which was composed of 139 genes including 116 source proteins, 7 signaling proteins and 16 TFs (which include NFKB2). Enrichment analysis for genes in this subnetwork identified several immune related categories and a few specific categories including: “SARS-CoV-2 innate immunity Evasion and Cell-Specific immune response (p-value=3.41e-4)” and “Hijack of Ubiquitination by SARS-CoV-2”.

**Figure 3:**
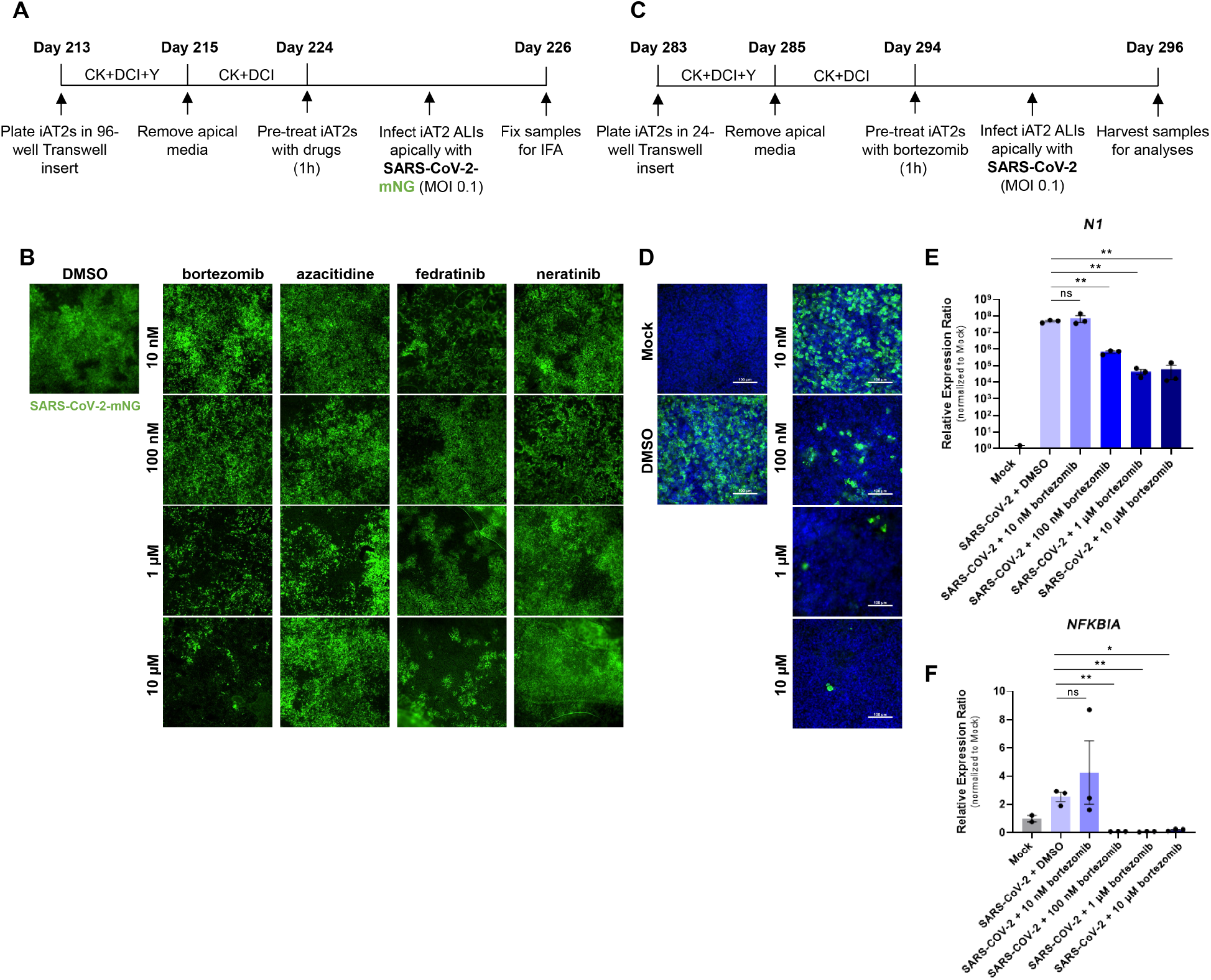
Validation of SDREM-based predicted genes and associated drugs in SARS-CoV-2-infected iAT2s. (A) Schematic of experimental setup for the screen of FDA-approved drugs on SARS-CoV-2 infection of iAT2s. (B) Representative fluorescent images (10x) of SARS-CoV-2-mNG-infected iAT2s pre-treated with four FDA-approved drugs that target the five proteins identified by SDREM (Table 2). (C) Schematic of experimental approach for bortezomib testing in iAT2s. (D) Representative immunofluorescence micrographs of SARS-CoV-2-infected iAT2s treated with bortezomib (10 nM-10 *μ*M) vs carrier vehicle (DMSO). DAPI (blue) and SARS-CoV-2 nucleocapsid protein (green) shown, scale bar = 100 *μ*m. (E, F) RT-qPCR of nucleocapsid N1 transcript and NF-kB pathway target gene NFKBIA in mock-infected or SARS-CoV-2-infected iAT2s treated with varying bortezomib concentrations (10 nM-10 *μ*M). \**p* < 0.05, * * *p* < 0.001, pairwise multiple comparisons. All bars represent mean ± standard deviation.

**Figure 4:**
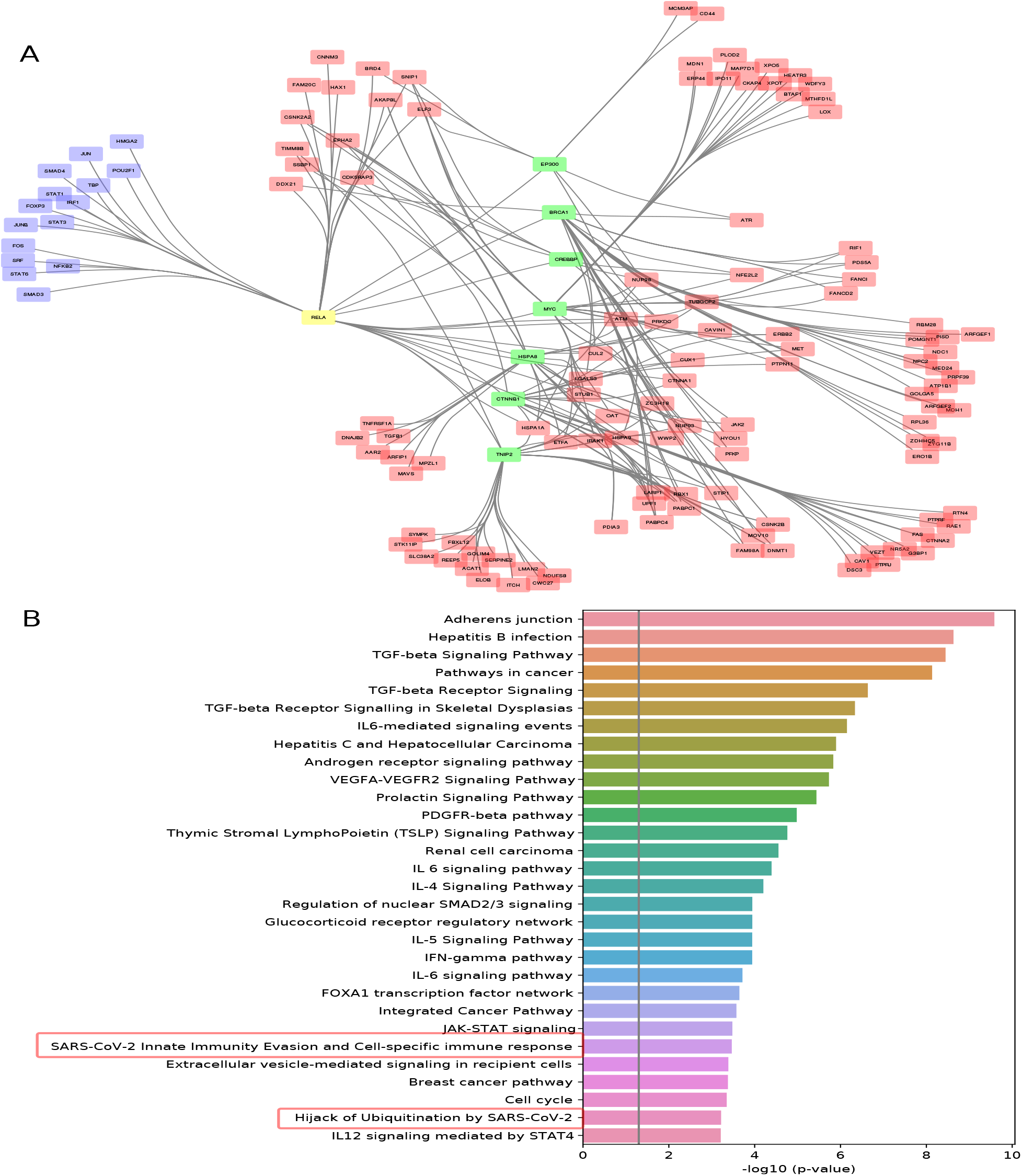
RELA / NFKB sub-network and enriched Pathways. (A) SDREM reconstructed RELA (yellow node) sub-network that connects direct targets of SARS-COV-2 (red) to TFs (blue) via signaling proteins (yellow and green). (B) Enriched Pahtwyays for genes in the RELA sub-network.

## Discussion

By integrating data from several relevant molecular resources, we were able to identify a subset of genes that are (1) connected to viral proteins in signaling pathways (2) impact downstream expression response to the infection and (3) identified to be DE in underlying conditions. We used computational methods that combine probabilistic graphical models with combinatorial network analysis to rank top genes and pairs of genes and to intersect these with underlying condition genes. Our methods identified a list of 19 genes in the overlap of all relevant datasets when looking at top single node rankings and 39 genes when looking at pair rankings. Functional analysis of these genes indicated that many are related to host response to viral infections and to replications, the two key types of pathways expected to be activated following infection. While some of the top ranked proteins are well known, several are novel predictions that have not been previously studied since they do not directly interact with SARS-CoV-2 proteins. As shown in Supplementary Table S4, 34 of the proteins in our intersection sets have known potential treatments, including 9 proteins associated with approved drugs (EGFR, ERBB2, ESRRA, HIF1A, IRAK1, JAK2, NR3C1, PLAU, RELA/NFKB, RORA and RXRA). Of these, 7 are not characterized as directly interacting with SARS-CoV-2 proteins.

We experimentally tested FDA-approved drugs that are associated with some of the predicted SDREM proteins. For these tests, we utilized a physiologically relevant human in vitro model system based on iPSC-derived alveolar epithelial type II cells (iAT2s), which we have previously shown to be similar to primary adult alveolar epithelial type II cells [27] and were recently established as a novel model for SARS-CoV-2 infections [13]. Of the four drugs we tested, bortezomib - a compound that modulates NF-kB activity by proteosomal inhibition - significantly inhibited SARS-CoV-2, with transcript data suggesting that bortezomib suppresses NF-kB activity in iAT2s. While these studies show the potential for the methods presented to be used to identify potential treatments for COVID-19, follow up studies are obviously needed to investigate the role of NF-kB inhibition on infection severity.

While our results help to support the computationally predicted hypothesis that downregulation of NF-kB pathway activity inhibits viral infection, there are several important caveats to this interpretation. First bortezomib does not act as a specific NF-kB pathway inhibitor, but rather acts through the more general mechanism of inhibition of the proteosome, with broad effects on cellular protein processing and degradation, beyond simply blocking the degradation of the NF-KB inhibitory protein, IKBa, which inhibits NF-*κ*B activity [28]. Indeed, the SARS-CoV-2 viral life cycle itself depends on cellular protein processing and so it is possible that proteosome inhibition with bortezomib may directly inhibit the viral life cycle [29, 18]. Furthermore, there is some controversy raised by some past published reports that found, in more prolonged treatment regimens for multiple myeloma, bortezomib treatment can in some circumstances activate NF-*κ*B activity in myeloma cells [30]. Thus, future studies using genetic knockdown of REL A/p656 and animal models studies of treating SARS-Co-V2 infection with bortezomib are needed to establish the usefulness of this treatment and its mode of action.

While our focus here was on SARS-Cov-2 infection the methods we developed and used are general and can be applied to model any viral infection. These methods complement experimental methods enabling large scale integration of time series omics data to identify key proteins in the host response pathways.

## Methods

### Datasets

We used both condition specific and general interaction data for learning dynamic viral infection models. We also collected lung expression data for conditions that were reported to impact SARS-CoV-2 mortality and infection rates. Below we provide information about all data used in this study.

### Viral host interactions and phosphorylation data

We used the SARS-CoV-2 and human protein-protein interactome reported by Gordon *et al*. [3] and Stukalov *et al*. [4] which identified 1396 protein interactions between 31 viral proteins and 1148 human proteins using affinity purification-mass spectrometry analysis. While this data is virus specific, we note that prior studies for other viruses indicated that a single screen is unlikely to fully cover the entire set of virus-host interactions [31]. Supplementary Table S5 provides the full list of interactions we used. We also used a protein phosphorylation data dataset in which phosphorylation levels were profiled at 0, 2, 4, 8, 12 and 24 hours post SARS-CoV-2 infection [18]. Using this data, we obtained 543 significantly phosphorylated proteins across all time points profiled (student t-test p-value<0.05 and log2fold change>0.4). Please refer to Supplementary Table S6 for the detailed list of phosphorylated proteins.

### RNAi or CRISPR knockdown data

We searched the literature for a list of RNAi or CRISPR screen experiments which test the impact of gene knockdown/knockout on coronavirus load. In particular, we collected RNAi screening data for 5 different coronaviruses: IBV-CoV, MERS-CoV, MHV-CoV, SARS-CoV and SARS-CoV-2 [32, 33, 34, 35, 36, 37, 38, 39, 40, 41]. Additionally, we collected CRISPR screens for SARS-CoV-2 [22, 23, 24, 42]. The combined set of RNAi and CRISPR screen hits is comprised of 535 human proteins in cells infected with a coronavirus, 40 of which were present in our initial SARS-CoV-2 and human network. Table 3 summarizes the hits used in this study while Supplementary Table S7 provides the full list of screen hits used in this study.

**Table 3:** Summary of literature-derived coronaviruses genome-wide RNA or CRISPR screens. For each screen, we list associated coronavirus, number of significant screen hits and corresponding references. We use 14 different RNAi or CRISPR screen studies for avian infectious bronchitis virus (IBV-CoV), Middle East respiratory syndrome (MERS-CoV), murine hepatitis virus (MHV-CoV) and severe acute respiratory syndrome (SARS-CoV and SARS-CoV-2) coronaviruses.

| Coronavirus | Screen hits | References |
| --- | --- | --- |
| IBV-CoV | 94 | [34, 37] |
| MERS-CoV | 20 | [38, 41] |
| MHV-CoV | 26 | [32, 33, 36] |
| SARS-CoV | 135 | [35, 39, 40] |
| SARS-CoV-2 | 272 | [22, 23, 24, 42] |

### General Protein-protein and protein-DNA interactions

Protein-DNA interactions were obtained from our previous work [43], which contains 59578 protein-DNA interactions for 399 Transcription Factors (TFs). Protein-protein interactions (PPIs) were obtained from the HIPPIE database [44], which contains more than 270000 annotated PPIs and for each provides a confidence score which was further used in our network analysis (see below).

### Time series transcriptomics

We used several longitudinal transcriptomics datasets to learn the regulatory and signaling networks underlying the SARS-CoV-2 infection. The first was a bulk expression dataset in which iPSC derived lung epithelial cells were infected and profiled before and 1 and 4 hours following infection [13]. The expression of SARS-CoV-2 viral proteins were also quantified for this dataset and we were able to obtain expression levels for 11 viral proteins which were further used in our analysis (see below).

In addition to the bulk data we also used a single-cell RNA-seq data on Calu-3 cell line profiled at 0 (mock), 4, 8, 12 hours post SARS-CoV-2 infection. We filtered all the cells with less than 200 expressed genes or with over 40% mitochondrial genes [14]. We also filtered expressed in less than 3 cells or with a very low dispersion (< 0.15).

We also applied our model to another time-series single-cell RNA-seq data on differentiated human bronchial epithelial cells at 0 (mock), 1, 2, 3 days post SARS-Cov-2 infection [15]. The low quality cells were filtered using the same method as for the Calu-3 data above.

### Underlying condition lung expression data

We collected and analyzed lung expression data from several reported underlying conditions that impact SARS-CoV-2 infection. Most of the data we used was from bulk microarray expression studies of lung tissues. These included studies focused on lung cancer [45] (Accession number: GSE2514), hypertension [19] (GSE24988), Diabetes [46] (GSE15900), COPD [47] (GSE38974), and smoking [48] (GSE10072). We also used single cell RNA-seq lung data for gender expression analysis and for inferring aging related genes from [20] (GSE136831). For each dataset, the differentially expressed genes are extracted using the R package ‘limma’ for microarray data [49] and by using a ranksum test for single cell data.

### Reconstructing dynamic signaling and regulatory networks using SDREM

For the analysis and modeling of SARS-CoV-2 infection in lung cells, we extend the Signaling Dynamic Regulatory Events Miner (SDREM) [12] method. SDREM integrates time-series bulk gene expression data with static PPIs and protein-DNA interaction to reconstruct response regulatory networks and signaling pathways. SDREM iterates between two methods. The first, DREM [50] uses an input-output hidden Markov model (IOHMM) to reconstruct dynamic regulatory networks by identifying bifurcation events where a set of co-expressed genes diverges. DREM annotates these splits and paths (co-expressed genes) with TFs that regulate genes in the outgoing upward and/or downward paths.

To extend the SDREM to the single-cell level, instead of using DREM as for analyzing bulk transcriptomic data, here we incorporated our previously developed methods SCDIFF and CSHMM [43, 51] to reconstruct the regulatory network underlying the single-cell time-series RNA-seq data. Based on the single-cell regulatory network inference, we generate a list of transcription factors with 3 metrics to evaluate their importance. First, percentage of regulated cells; a TF that regulates a higher percentage of cells will also be assigned with an importance score. Second, p-value for the TF regulation; we calculated a hypergeometric test p-value for each of the predicted TFs based on their target genes. If the target genes of a TF are differentially expressed between points that the TF is regulating, such TF-regulation would be considered more reliable and it should be weighted with a higher score. Namely, TFs will a smaller p-value will be assigned with a higher importance score. Third, we also evaluate the TFs based on their Lasso logistic coefficients from the single-cell regulatory inference (by SCDIFF); a TF with a higher coefficient should be treated with a higher importance. Finally, we calculate an overall importance score *IS* for each of the TFs predicted from the single-cell regulatory network inference method based on the above 3 scoring metrics.

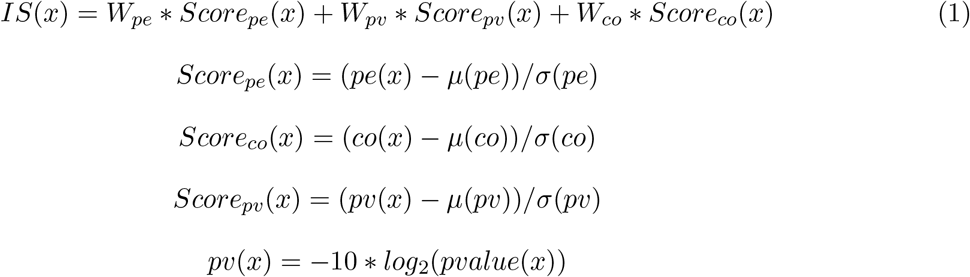

Where *pe*(*x*), *co*(*x*), *pvalue*(*x*) are the percentage score, coefficient score, and p-value score for TF *x* respectively. *W*_*pe*_, *W*_*pv*_, *W*_*co*_ are weights for each of the scoring metrics. By default, they are all set as 1 for equal importance. However, users are allowed to specify a different set of weights to emphasize specific scoring metrics. If a TF is found to regulate multiple edges of the reconstructed trajectory and thus have multiple overall scores, the maximal one will be used. We ranked all the predicted TFs based on their overall scores. The top ones (under a specific cutoff parameter specified by SDREM method) will be chosen as the final predicted regulators for the following analyses.

The second part of SDREM uses a network orientation method [11], which orients the undirected protein interaction edges such that the targets can be explained by relatively short, high-confidence pathways that originated at the inputs with provided PPIs, source proteins, and target TFs. Generally, SDREM searches for high scoring paths that start at the virus proteins, continues with the host proteins they interact with and ends with TFs and their targets. By iterating between identifying TFs based on the expression of their targets and connecting identified TFs to source (virus) proteins, SDREM can identify a set of high scoring pathways and regulators. These are then analyzed using graph-based scoring methods as we discuss below to identify key proteins mediating viral signals.

### Using human SARS-CoV-2 transcriptomics and protein phosphorylation data with SDREM

We modified SDREM to improve its performance on the SARS-CoV-2 transcriptomics data. To remove the potential batch effect, we have performed cross-sample normalization between the bulk RNA-seq sample between different time points (control, 1dpi, 4dpi). Another extension we applied is the use of the viral expression data to remove sources and their connected host proteins. Specifically, we first identified the non-expressed viral genes in those datasets. Then, we remove all the host source proteins that interact with the viral proteins that correspond to those non-expression viral genes. We integrate the protein phosphorylation data by adjusting the prior for significantly phosphorylated proteins. We first set the prior of all proteins as a default value (e.g, 0.5). Then we scaled the log2fc of the significantly phosphorylated proteins to the range of [0.5, 1] with a min-max normalization method. To mitigate the impact of the outliers, we replaced the min-max values with 5% and 95% percentiles. Proteins (nodes) that are highly phosphorylated will be assigned with a larger prior and they will be favored in the SDREM network analysis.

### Identifying key genes in SDREM reconstructed networks

To rank top genes identified by SDREM, we used the strategy described in [11] to estimate *in silico* effects of removing a protein from the signaling network component of an SDREM model. The method computes how the connectivity to the TFs is affected when a node (gene) or a combination of nodes, is removed. Intuitively, this score captures the impact of the removal on the path weights that remain for linking TFs to sources (eqn.2).

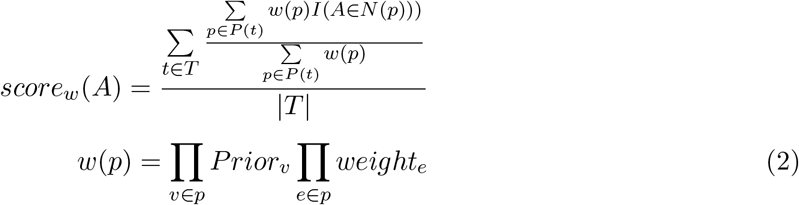

Where *A* is the deleted node, *T* is the set of all targets, *P*(*t*) is the set of paths to the target *t* to be considered, *I*(*) is an indicator function that has the value 1 if the condition * is satisfied, *N* (*p*) is the set of nodes on the path *p,w*(*p*) is the path weight, which is the product of all node priors *Prior*_*v*_ and edge weights *weight*_*e*_ in the path. The node prior is determined using the protein phosphorylation data (if available). The edge weight is based on the strength for the interaction (e.g., PPI) between the source and target nodes of the edge.

Although single gene inference might be very informative, higher-order knockdowns (of two or more genes) may prove to be more robust because they can target several pathways simultaneously. Experimentally testing all possible gene combinations would be prohibitive but in-silico analysis is much faster given the relatively small number of genes in the resulting SDREM network. Scores for pair removals are computed in a similar way to individual scores, by finding double knockdown has a stronger effect than expected based on the score for individual gene of the pair, which corresponds to lower value of 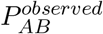 (see eqn.3 for details).

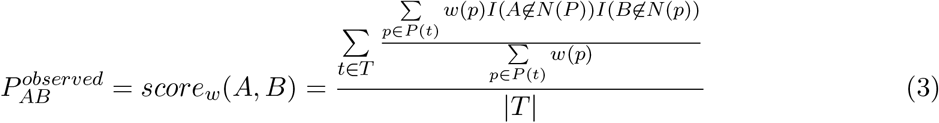

The above eqn.3 denotes the average fraction of path weight that remains after removing paths that contain node *A* and *B*.

Several rankings can be derived based on the score computed above. These differ in the paths used for the scoring (top ranked or all), weather target connectivity is evaluated separately for every source or for all sources combined and using weighted versus unweighted versions of the SDREM network. See Supplementary Table S1 (Meta) for details on what was used in the analysis.

### Analyzing underlying condition data

We examined several recent studies to determine underlying condition that impact SARS-CoV-2 mortality and infection rates [7, 8]. Based on these we selected seven different conditions for which we were able to obtain lung expression data (Supplementary Methods). We next used the ranksum statistical test to compute DE p-value for each genes in each condition. Finally, we used the assigned p-values and expression fold change signs to compute quantile value between 0 and 1 for each gene in each condition and divided genes in top quantile for each condition to ‘over-expressed’ and ‘repressed’. We next intersected the list of top ranked genes from the SDREM reconstructed networks with genes identified as significantly up or down regulated in the underlying conditions data. We identified several parameters for which the intersection is significant meaning that many of the top SDREM genes are also DE in several of the conditions. See Supplementary Figures S3 and S4 for SDREM result, Supplementary Figures S5 and S6 for bulk SDREM result with phosphorylation, Supplementary Figures S7 and S8 for the first single-cell SDREM result with phosphorylation, and Supplementary Figures S9 and S10 for the second single-cell SDREM result.

### Potential treatments for top genes

In a similar fashion to Gordon *et al*. [3], we searched public resources (ChEMBL25 [52], IUPHAR/BPS, Pharos [53] and ZINC [54]), as well as literature in order to identify existing drugs and reagents that directly modulate the candidate genes derived from our network reconstruction and condition-specific analyses. Supplementary Table S4 provides a full list of drugs and reagents targeting the identified candidate genes whereas Table 2 provides a list of approved drugs targeting five highly-ranked genes.

### SARS-CoV-2 infection

SARS-CoV-2 isolate USA WA1/2020 (GenBank Accession number MN985325) was kindly provided by CDC’s Principal Investigator Natalie Thornburg and the World Reference Center for Emerging Viruses and Arboviruses (WRCEVA). Recombinant SARS-CoV-2 expressing mNeonGreen (SARS-CoV-2-mNG) was kindly provided by Pey-Yong Shi, University of Texas Medical Branch, Galveston and the WRCEVA [26]. This virus is based on SARS-CoV-2 isolate USA WA1/2020. SARS-CoV-2 and SARS-CoV-2-mNG stocks were grown in Vero E6 cells and virus titers were determined by tissue culture infectious dose 50 (TCID50) assays as described before by Huang *et al*. [13]. All work with SARS-CoV-2 and SARS-CoV-2-mNG was performed in the BSL-4 facility of the National Emerging Infectious Diseases Laboratories (NEIDL) at Boston University following approved SOPs.

### Treatment and infection of cells

Human induced pluripotent stem cell-derived alveolar type II cells (iAT2s) were generated as previously described via directed differentiation [27, 55], and the resulting putative iAT2s (¿200 days of differentiation) were maintained in 3D Matrigel in CKDCI media. For SARS-CoV-2 infections, air-liquid interface cultures of iAT2s were prepared by seeding iAT2s in 24-well or 96-well transwells coated with Matrigel (Corning, Cat No. 354277) in CKDCI+Y-27632, removing the apical media after 48 hours, and feeding with CKDCI every 2 days [13]. For the initial drug testing, azacitidine (Millipore Sigma, Cat No. A2385), bortezomib (Selleckchem, Cat No. S1013), fedratinib (Selleckchem, Cat No. S2736), and neratinib (Selleckchem, Cat No. S2150) were dissolved in DMSO and diluted in CKDCI media at the indicated concentrations and added apically and basolaterally to iAT2s seeded in 96-well transwells (Corning, Cat No. CLS3381) at 80,000 cells/well. After 1 hour of pre-treatment, the apical treatments were removed and cells were mock-infected or infected with SARS-CoV-2-mNG at a multiplicity of infection (MOI) of 0.1. Diluted drug treatments were left basolaterally for the entirety of the infection. Inocula were removed 1 hour post-infection, returning iAT2s to ALI. Two days post-infection, the cells were fixed in 10% neutral buffered formalin for at least 6 hours at 4°*C* and removed from the BSL-4 laboratory. Follow-up testing with bortezomib was performed as above but with the following differences: iAT2s seeded in 24-well transwells (300,000 cells/well) were used and these infections were performed using SARS-CoV-2 at a MOI of 0.1.

### Immunostaining and microscopic analysis

For antibody staining, the cells were permeabilized with 1:1 (vol:vol) acetone-methanol solution for 5 minutes at -20°*C*, incubated in 0.1 M glycine for 10 minutes at room temperature, and subsequently incubated in blocking reagent (2% bovine serum albumin, 0.2% Tween 20, 3% glycerin, and 0.05% NaN3 in PBS) for 15 minutes at room temperature. After each step, the cells were washed three times in PBS. The cells were incubated for one hour at room temperature with a rabbit antibody directed against the SARS-CoV nucleocapsid protein (Rockland; 100 *μ*l per well, diluted 1:2000 in blocking reagent; this antibody cross-reacts with the SARS-CoV-2 nucleocapsid protein). The cells were washed three times in PBS and incubated with the secondary antibody for 1 hour at room temperature (goat anti-rabbit antibody conjugated with AlexaFluor488 (Invitrogen); diluted 1:200 in blocking reagent) and 4’,6-diamidino-2-phenylindole (DAPI) (Sigma-Aldrich) was used at 200 ng/mL for nuclei staining. For cells infected with SARS-CoV-2-mNG, only the DAPI stain step was performed. Membranes were excised from the transwell supports and mounted on slides using FluorSave mounting medium (Millipore) and glass coverslips, and slides were stored at 4°*C* prior to imaging. Images were acquired at 4x, 10x, and 30x magnification using a Nikon Eclipse Ti2 microscope with Photometrics Prime BSI camera and NIS Elements AR software.

### Software and data availability

The single-cell extension of the SDREM model (named scSDREM) is implemented in Python. It’s publicly available at https://github.com/phoenixding/sdremsc. The bulk and single-cell time-series SARS-CoV-2 viral infection transcriptomics data used in this work are available under the GEO accession number GSE153277 and GSE148729, respectively. The regulatory models (by iDREM) for the bulk and single-cell SARS-CoV-2 transcriptomics data is available at : http://www.cs.cmu.edu/~jund/sars-cov-2

## Acknowledgements

The authors thank Natalie Thornburg, CDC, Pey-Yong Shi, UTMB Galveston, and the World Reference Center for Emerging Viruses and Arboviruses for sharing viruses. This work was partially supported by NIH grant 1R01GM122096 and by a C3.ai DTI Research Award to ZB-J; NIH grants NO1 75N92020C00005, U01TR001810, and an Evergrande COVID-19 Response Fund Award from the Massachusetts Consortium on Pathogen Readiness (MassCPR) to DNK. EM acknowledges SARS-CoV-2 funding from Fast Grants, Evergrande MassCPR, and NIH NCATS grant UL1TR001430. ELS is supported by NIH HL007035T32 training grant Biology of the Lung.

## Author contributions

ZB-J, JD, JL-M, DNK, EM, and YY designed the research; JD, JL-M, and YY developed and implemented the methods ; JH prepared iPSC-derived iAT2s and performed RT-PCR; AJH performed the drug treatment and infection studies; ELS performed immunofluorescence analysis and microscopy; JD, JL-M and YY analyzed data ; and JD, JL-M, YY, JH, AJH, ELS, EM, DNK, and ZB-J wrote the manuscript.

## Conflict of interest

None.

## Notes

### Competing Interest Statement

The authors have declared no competing interest.

### Summary of Updates

- Updated to include recent genome-wide CRISPR screens for SARS-CoV-2. - Experimentally tested treatments for a number of the predicted targets. - Showed that blocking one of the predicted indirect targets significantly reduces viral loads in stem cell-derived alveolar epithelial type II cells (iAT2s).

