## Supplementary figures and images for "Reconstructed signaling and regulatory networks identify potential drugs for SARS-CoV-2 infection"

### Supplementary Figure S1

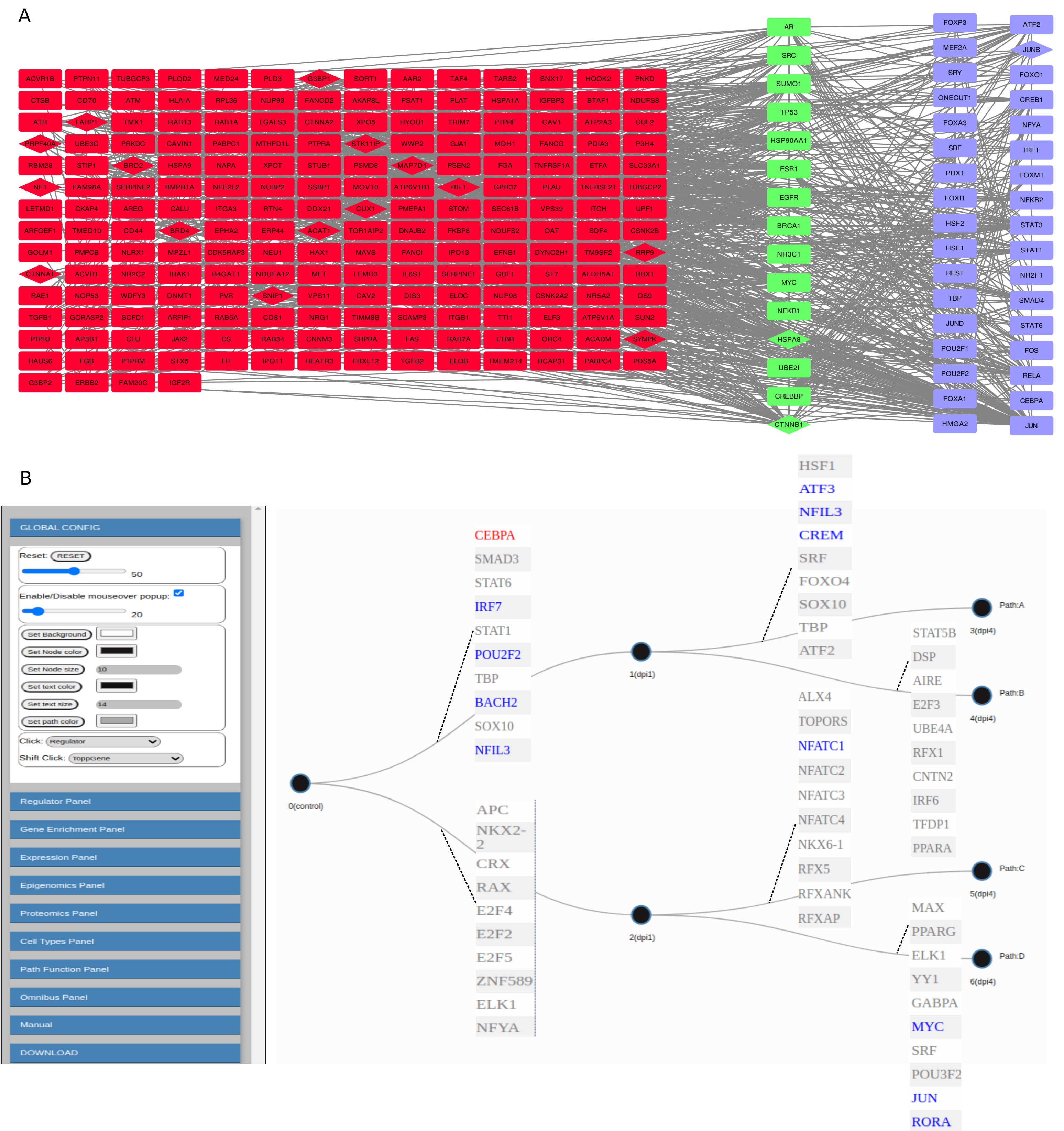

### Supplementary Figure S2

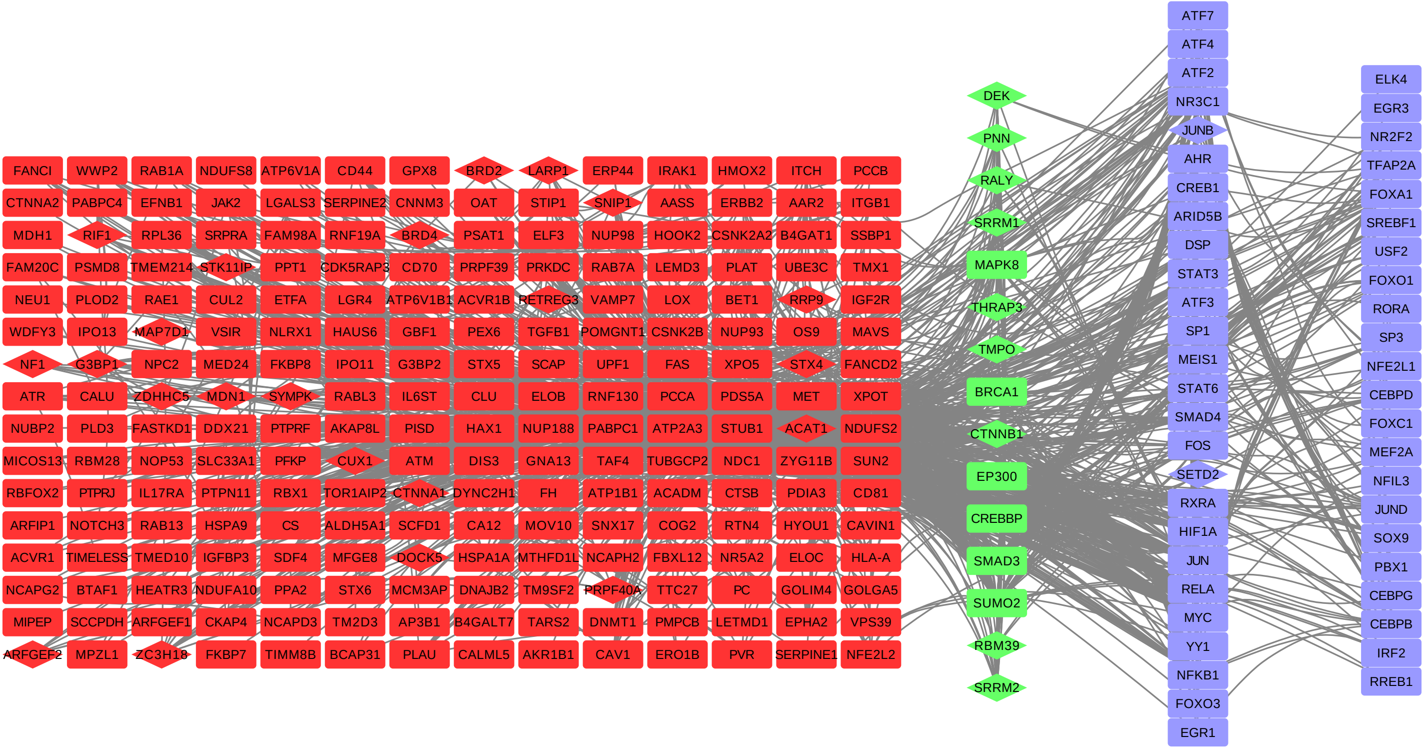
