## Supplementary Figure S3 for "Reconstructed signaling and regulatory networks identify potential drugs for SARS-CoV-2 infection"

**A**

Sum threshold

| quantile | 0 | 1 | 2 | 3 | 4 | 5 | 6 |
|---|---|---|---|---|---|---|---|
| 0.01 | 0.535492 | 2.08E-05 | 0.106374 | 1 | 1 | 1 | 1 |
| 0.05 | 0.535492 | 0.000922 | 0.008137 | 0.169801 | 1 | 1 | 1 |
| 0.1 | 0.535492 | 0.00142 | 0.000842 | 0.0014 | 0.007085 | 0.014895 | 1 |
| 0.15 | 0.535492 | 0.016288 | 4.96E-06 | 0.000375 | 0.050248 | 0.053524 | 1 |
| 0.2 | 0.535492 | 0.03398 | 9.33E-06 | 9.59E-05 | 0.145524 | 0.118562 | 1 |
| 0.25 | 0.535492 | 0.037615 | 2.63E-06 | 6.50E-06 | 0.290212 | 1 | 1 |
| 0.3 | 0.535492 | 0.129611 | 0.000583 | 7.22E-05 | 0.174307 | 1 | 1 |

**B**

Sum threshold

| quantile | 0 | 1 | 2 | 3 | 4 | 5 | 6 |
|---|---|---|---|---|---|---|---|
| 0.01 | 63 | 16 | 1 | 0 | 0 | 0 | 0 |
| 0.05 | 63 | 33 | 6 | 1 | 0 | 0 | 0 |
| 0.1 | 63 | 45 | 13 | 5 | 2 | 1 | 0 |
| 0.15 | 63 | 46 | 23 | 8 | 2 | 1 | 0 |
| 0.2 | 63 | 49 | 27 | 11 | 2 | 1 | 0 |
| 0.25 | 63 | 52 | 33 | 15 | 2 | 0 | 0 |
| 0.3 | 63 | 49 | 30 | 15 | 3 | 0 | 0 |

**C**

| gene_symbol | hypertension | COPD | diabetes | smoke | cancer | sex | age | sum |
|---|---|---|---|---|---|---|---|---|
| cav1 | -1 | -1 | -1 | -1 | -1 | 1 | 0 | -4 |
| jun | -1 | 1 | -1 | -1 | -1 | -1 | 0 | -4 |
| hspa8 | -1 | -1 | 0 | 0 | 0 | -1 | 0 | -3 |
| erbb2 | 1 | -1 | 0 | -1 | -1 | 0 | -1 | -3 |
| srf | -1 | 1 | -1 | -1 | -1 | 0 | 0 | -3 |
| fos | -1 | 0 | 1 | -1 | -1 | 0 | -1 | -3 |
| acat1 | 0 | -1 | -1 | -1 | 0 | -1 | 1 | -3 |
| ctnna1 | 1 | -1 | 0 | -1 | -1 | -1 | 0 | -3 |
| brd2 | -1 | 1 | -1 | -1 | -1 | -1 | 1 | -3 |
| myc | -1 | 1 | 0 | 0 | 0 | -1 | -1 | -2 |
| akap8l | 0 | -1 | -1 | 0 | 0 | -1 | 1 | -2 |
| stx4 | -1 | -1 | 0 | 0 | 0 | 0 | 0 | -2 |
| creb1 | 0 | -1 | 1 | 1 | -1 | -1 | -1 | -2 |
| junb | -1 | 1 | 0 | 0 | -1 | -1 | 0 | -2 |
| crebbp | 0 | -1 | -1 | -1 | 0 | 0 | 1 | -2 |
| irf1 | -1 | 0 | 1 | -1 | -1 | 1 | -1 | -2 |
| stat6 | 0 | 0 | -1 | -1 | -1 | 0 | 1 | -2 |
| rela | -1 | 0 | 0 | 0 | -1 | 1 | -1 | -2 |
| stip1 | -1 | 1 | 1 | 1 | 0 | 0 | 0 | 2 |
| cul2 | 0 | 1 | 0 | 0 | 0 | 0 | 1 | 2 |
| atm | 1 | 1 | 0 | 1 | -1 | 0 | 0 | 2 |
| g3bp1 | 0 | 0 | -1 | 1 | 1 | 0 | 1 | 2 |
| arfgef2 | 1 | 0 | -1 | 1 | 1 | -1 | 1 | 2 |
| prpf40a | 0 | 0 | 0 | 1 | 1 | -1 | 1 | 2 |
| bclaf1 | 0 | -1 | 1 | 1 | 0 | 1 | 0 | 2 |
| smad3 | 0 | -1 | 1 | -1 | 1 | 1 | 1 | 2 |
| larp1 | 0 | 0 | 1 | -1 | 1 | 0 | 1 | 2 |
| sympk | 0 | 0 | 0 | 0 | 1 | 1 | 1 | 3 |
| ep300 | 0 | 1 | 1 | -1 | 0 | 1 | 1 | 3 |
| ptprj | -1 | 1 | 1 | 0 | 0 | 1 | 1 | 3 |
| csnk2a2 | 0 | 1 | 1 | -1 | 1 | 0 | 1 | 3 |
| zc3h18 | 0 | 1 | 0 | 0 | 0 | 1 | 1 | 3 |
| itch | 0 | 1 | 0 | 1 | 0 | 0 | 1 | 3 |

**D**

| GO biological process complete | raw P value | ▲ FDR |
|---|---|---|
| positive regulation of gene expression | 1.93E-12 | 3.07E-08 |
| positive regulation of nitrogen compound metabolic process | 1.07E-11 | 5.66E-08 |
| cellular response to chemical stimulus | 1.80E-11 | 5.74E-08 |
| response to stress | 1.46E-11 | 5.81E-08 |
| positive regulation of macromolecule metabolic process | 8.77E-12 | 6.97E-08 |
| cellular response to organic substance | 3.54E-11 | 8.04E-08 |
| positive regulation of cellular metabolic process | 3.42E-11 | 9.05E-08 |
| positive regulation of metabolic process | 5.23E-11 | 9.24E-08 |
| symbiotic process | 5.91E-11 | 9.39E-08 |
| gland development | 5.00E-11 | 9.92E-08 |
| positive regulation of macromolecule biosynthetic process | 8.96E-11 | 1.29E-07 |
| viral process | 2.37E-10 | 2.51E-07 |
| positive regulation of cellular biosynthetic process | 2.08E-10 | 2.54E-07 |
| positive regulation of RNA metabolic process | 2.29E-10 | 2.60E-07 |
| positive regulation of transcription by RNA polymerase II | 2.03E-10 | 2.69E-07 |
| positive regulation of biosynthetic process | 2.72E-10 | 2.70E-07 |
| response to organic substance | 3.50E-10 | 3.27E-07 |
| positive regulation of cellular process | 3.75E-10 | 3.31E-07 |
| positive regulation of biological process | 4.41E-10 | 3.69E-07 |
| positive regulation of transcription, DNA-templated | 4.88E-10 | 3.88E-07 |
| regulation of cellular macromolecule biosynthetic process | 7.55E-10 | 5.72E-07 |
| response to chemical | 9.30E-10 | 6.72E-07 |
