## Supplementary Figure S4 for "Reconstructed signaling and regulatory networks identify potential drugs for SARS-CoV-2 infection"

**A**

Sum threshold (columns) / quantile (rows)

| quantile | 0 | 1 | 2 | 3 | 4 | 5 | 6 |
|---|---|---|---|---|---|---|---|
| 0.01 | 0.529557 | 1.29E-06 | 0.149318 | 1 | 1 | 1 | 1 |
| 0.05 | 0.529557 | 3.74E-05 | 0.003766 | 0.234769 | 1 | 1 | 1 |
| 0.1 | 0.529557 | 0.000337 | 5.28E-05 | 0.001174 | 0.014072 | 0.021348 | 1 |
| 0.15 | 0.529557 | 0.021332 | 2.95E-06 | 0.000213 | 0.09362 | 0.07605 | 1 |
| 0.2 | 0.529557 | 0.012269 | 6.39E-07 | 0.000491 | 0.074106 | 0.165951 | 1 |
| 0.25 | 0.529557 | 0.015355 | 1.59E-07 | 1.57E-07 | 0.45411 | 1 | 1 |
| 0.3 | 0.529557 | 0.094155 | 3.90E-06 | 4.05E-05 | 0.336754 | 1 | 1 |

**B**

Sum threshold (columns) / quantile (rows)

| quantile | 0 | 1 | 2 | 3 | 4 | 5 | 6 |
|---|---|---|---|---|---|---|---|
| 0.01 | 91 | 22 | 1 | 0 | 0 | 0 | 0 |
| 0.05 | 91 | 49 | 8 | 1 | 0 | 0 | 0 |
| 0.1 | 91 | 63 | 19 | 6 | 2 | 1 | 0 |
| 0.15 | 91 | 61 | 29 | 10 | 2 | 1 | 0 |
| 0.2 | 91 | 71 | 37 | 12 | 3 | 1 | 0 |
| 0.25 | 91 | 75 | 45 | 21 | 2 | 0 | 0 |
| 0.3 | 91 | 70 | 47 | 19 | 3 | 0 | 0 |

**C**

| gene_symbol | hypertension | COPD | diabetes | smoke | cancer | sex | age | sum |
|---|---|---|---|---|---|---|---|---|
| cav1 | -1 | -1 | -1 | -1 | -1 | 1 | 0 | -4 |
| jun | -1 | 1 | -1 | -1 | -1 | -1 | 0 | -4 |
| hspa8 | -1 | -1 | 0 | 0 | 0 | -1 | 0 | -3 |
| hax1 | -1 | -1 | 0 | -1 | 1 | -1 | 0 | -3 |
| erbb2 | 1 | -1 | 0 | -1 | -1 | 0 | -1 | -3 |
| srf | -1 | 1 | -1 | -1 | -1 | 0 | 0 | -3 |
| fos | -1 | 0 | 1 | -1 | -1 | 0 | -1 | -3 |
| acat1 | 0 | -1 | -1 | -1 | 0 | -1 | 1 | -3 |
| ctnna1 | 1 | -1 | 0 | -1 | -1 | -1 | 0 | -3 |
| brd2 | -1 | 1 | -1 | -1 | -1 | -1 | 1 | -3 |
| sympk | 0 | 0 | 0 | 0 | 1 | 1 | 1 | 3 |
| ep300 | 0 | 1 | 1 | -1 | 0 | 1 | 1 | 3 |
| rab1a | 0 | 1 | 1 | -1 | 1 | 0 | 1 | 3 |
| irak1 | 0 | 0 | 1 | -1 | 1 | 1 | 1 | 3 |
| ptprj | -1 | 1 | 1 | 0 | 0 | 1 | 1 | 3 |
| neu1 | -1 | 1 | 1 | 1 | 1 | -1 | 1 | 3 |
| csnk2a2 | 0 | 1 | 1 | -1 | 1 | 0 | 1 | 3 |
| zc3h18 | 0 | 1 | 0 | 0 | 0 | 1 | 1 | 3 |
| fgb | -1 | 1 | 1 | 1 | 1 | 1 | -1 | 3 |
| plau | 0 | 0 | 0 | 1 | 1 | 1 | 0 | 3 |
| itch | 0 | 1 | 0 | 1 | 0 | 0 | 1 | 3 |

**D**

| GO biological process complete | raw P value | ▲ FDR |
|---|---|---|
| response to external stimulus | 3.73E-09 | 5.92E-05 |
| positive regulation of cellular process | 5.08E-08 | 4.04E-04 |
| positive regulation of nitrogen compound metabolic process | 9.34E-08 | 4.95E-04 |
| positive regulation of metabolic process | 1.27E-07 | 5.04E-04 |
| positive regulation of cellular metabolic process | 1.98E-07 | 6.31E-04 |
| positive regulation of macromolecule metabolic process | 3.81E-07 | 7.57E-04 |
| positive regulation of biological process | 2.87E-07 | 7.62E-04 |
| regulation of DNA-binding transcription factor activity | 3.70E-07 | 8.40E-04 |
| response to stress | 5.21E-07 | 9.20E-04 |
| positive regulation of protein metabolic process | 6.62E-07 | 1.05E-03 |
| regulation of multicellular organismal process | 9.91E-07 | 1.43E-03 |
| response to endogenous stimulus | 1.19E-06 | 1.57E-03 |
| response to extracellular stimulus | 1.41E-06 | 1.72E-03 |
| response to hormone | 1.74E-06 | 1.73E-03 |
| regulation of protein metabolic process | 1.70E-06 | 1.80E-03 |
| regulation of protein modification process | 1.63E-06 | 1.85E-03 |
| negative regulation of programmed cell death | 4.13E-06 | 2.85E-03 |
| symbiotic process | 3.63E-06 | 2.89E-03 |
| positive regulation of protein modification process | 3.88E-06 | 2.94E-03 |
| regulation of signaling | 4.10E-06 | 2.96E-03 |
| negative regulation of apoptotic process | 3.60E-06 | 3.01E-03 |
| regulation of cell communication | 3.59E-06 | 3.17E-03 |
| response to organic substance | 3.48E-06 | 3.26E-03 |
| response to chemical | 6.57E-06 | 4.35E-03 |
