## Supplementary Figure S5 for "Reconstructed signaling and regulatory networks identify potential drugs for SARS-CoV-2 infection"

**A**

Sum threshold (columns) × quantile (rows)

| quantile | 0 | 1 | 2 | 3 | 4 | 5 | 6 |
|---|---|---|---|---|---|---|---|
| 0.01 | 0.525971 | 1.67E-05 | 0.188794 | 1 | 1 | 1 | 1 |
| 0.05 | 0.525971 | 6.91E-06 | 7.27E-06 | 0.292627 | 1 | 1 | 1 |
| 0.1 | 0.525971 | 0.000209 | 3.03E-06 | 2.56E-05 | 0.022711 | 0.027533 | 1 |
| 0.15 | 0.525971 | 0.00267 | 2.88E-08 | 2.48E-05 | 0.142231 | 0.097275 | 1 |
| 0.2 | 0.525971 | 0.012907 | 1.50E-08 | 2.46E-07 | 0.130738 | 0.209256 | 1 |
| 0.25 | 0.525971 | 0.01349 | 1.52E-07 | 5.66E-07 | 0.321113 | 1 | 1 |
| 0.3 | 0.525971 | 0.099369 | 0.000115 | 1.74E-05 | 0.276062 | 1 | 1 |

**B**

Sum threshold (columns) × quantile (rows)

| quantile | 0 | 1 | 2 | 3 | 4 | 5 | 6 |
|---|---|---|---|---|---|---|---|
| 0.01 | 118 | 23 | 1 | 0 | 0 | 0 | 0 |
| 0.05 | 118 | 62 | 14 | 1 | 0 | 0 | 0 |
| 0.1 | 118 | 78 | 25 | 9 | 2 | 1 | 0 |
| 0.15 | 118 | 84 | 39 | 13 | 2 | 1 | 0 |
| 0.2 | 118 | 88 | 48 | 20 | 3 | 1 | 0 |
| 0.25 | 118 | 94 | 53 | 23 | 3 | 0 | 0 |
| 0.3 | 118 | 88 | 50 | 23 | 4 | 0 | 0 |

**C**

| gene_symbol | hypertension | COPD | diabetes | smoke | cancer | sex | age | sum |
|---|---|---|---|---|---|---|---|---|
| cav1 | -1 | -1 | -1 | -1 | -1 | 0 | 0 | -5 |
| ube2i | 0 | -1 | -1 | -1 | -1 | 0 | 0 | -4 |
| jun | -1 | 1 | -1 | -1 | -1 | -1 | 0 | -4 |
| atp6v1a | -1 | -1 | -1 | 1 | -1 | 0 | 0 | -3 |
| irf1 | -1 | 0 | 1 | -1 | -1 | 0 | -1 | -3 |
| erbb2 | 1 | -1 | 0 | -1 | -1 | 0 | -1 | -3 |
| fos | -1 | 0 | 1 | -1 | -1 | 0 | -1 | -3 |
| raly | 0 | 0 | -1 | -1 | 0 | -1 | 0 | -3 |
| acat1 | 0 | -1 | -1 | 0 | 0 | -1 | 0 | -3 |
| hsf1 | -1 | 0 | 0 | -1 | 0 | -1 | 0 | -3 |
| ctnna1 | 0 | -1 | 0 | -1 | -1 | 0 | 0 | -3 |
| brd2 | -1 | 1 | -1 | -1 | -1 | -1 | 1 | -3 |
| nfe2l2 | -1 | -1 | 1 | 0 | -1 | 0 | 0 | -2 |
| dis3 | -1 | 0 | 0 | -1 | -1 | 0 | 1 | -2 |
| myc | -1 | 1 | 0 | 0 | 0 | -1 | -1 | -2 |
| akap8l | 0 | -1 | -1 | 0 | 0 | -1 | 1 | -2 |
| foxa3 | -1 | 0 | 0 | 0 | 0 | 0 | -1 | -2 |
| stx4 | -1 | -1 | 0 | 0 | 0 | 0 | 0 | -2 |
| junb | -1 | 1 | 0 | 0 | -1 | -1 | 0 | -2 |
| srf | -1 | 1 | 0 | -1 | -1 | 0 | 0 | -2 |
| sox12 | 0 | -1 | 0 | -1 | 0 | 0 | 0 | -2 |
| mef2a | 0 | 0 | 0 | -1 | -1 | 0 | 0 | -2 |
| stat6 | 0 | 0 | -1 | -1 | -1 | 0 | 1 | -2 |
| clu | 1 | -1 | -1 | -1 | -1 | 0 | 1 | -2 |
| nr2f1 | 0 | -1 | 0 | -1 | -1 | 0 | 1 | -2 |
| sox10 | 0 | 1 | 1 | 1 | -1 | 0 | 0 | 2 |
| sympk | 0 | 0 | 0 | 0 | 0 | 1 | 1 | 2 |
| ep300 | 0 | 0 | 1 | -1 | 0 | 1 | 1 | 2 |
| cul2 | 0 | 1 | 0 | 0 | 0 | 0 | 1 | 2 |
| atm | 1 | 1 | 0 | 1 | -1 | 0 | 0 | 2 |
| atr | 1 | 0 | 0 | 0 | 1 | 0 | 0 | 2 |
| g3bp1 | 0 | 0 | -1 | 1 | 1 | 0 | 1 | 2 |
| rif1 | 0 | 1 | 0 | 0 | -1 | 1 | 1 | 2 |
| ptprj | -1 | 1 | 1 | 0 | 0 | 1 | 0 | 2 |
| prpf40a | 0 | 0 | 0 | 1 | 1 | -1 | 1 | 2 |
| oat | 1 | 0 | -1 | 0 | 1 | 1 | 0 | 2 |
| rad18 | 1 | 0 | 0 | 0 | 0 | 0 | 1 | 2 |
| smad3 | 0 | -1 | 1 | -1 | 1 | 1 | 1 | 2 |
| foxm1 | 1 | 0 | 0 | 0 | 1 | 0 | 0 | 2 |
| larp1 | 0 | 0 | 1 | -1 | 1 | 0 | 1 | 2 |
| brca1 | 1 | 0 | 0 | 1 | 1 | 0 | 0 | 3 |
| onecut1 | 0 | 1 | -1 | 1 | 1 | 0 | 1 | 3 |
| sox2 | 1 | -1 | 1 | 1 | 1 | 0 | 0 | 3 |
| zc3h18 | 0 | 1 | 0 | 0 | 0 | 1 | 1 | 3 |
| plau | 0 | 0 | 0 | 1 | 1 | 1 | 0 | 3 |
| itch | 0 | 1 | 0 | 1 | 0 | 0 | 1 | 3 |
| csnk2b | 1 | 1 | 0 | 0 | 1 | 0 | 0 | 3 |
| pdia3 | 0 | 1 | -1 | 0 | 1 | 1 | 1 | 3 |

**D**

| GO biological process complete | raw P value | FDR |
|---|---|---|
| positive regulation of nitrogen compound metabolic process | 3.34E-17 | 5.31E-13 |
| positive regulation of cellular metabolic process | 2.03E-16 | 1.61E-12 |
| positive regulation of macromolecule metabolic process | 9.69E-16 | 5.13E-12 |
| positive regulation of cellular biosynthetic process | 4.28E-15 | 1.70E-11 |
| positive regulation of biosynthetic process | 6.49E-15 | 2.06E-11 |
| positive regulation of transcription by RNA polymerase II | 8.40E-15 | 2.23E-11 |
| positive regulation of metabolic process | 1.33E-14 | 2.64E-11 |
| positive regulation of macromolecule biosynthetic process | 1.22E-14 | 2.77E-11 |
| positive regulation of cellular process | 1.76E-14 | 3.10E-11 |
| positive regulation of nucleobase-containing compound metabolic process | 1.24E-13 | 1.79E-10 |
| positive regulation of transcription, DNA-templated | 1.18E-13 | 1.87E-10 |
| positive regulation of gene expression | 1.88E-13 | 2.49E-10 |
| positive regulation of RNA biosynthetic process | 3.90E-13 | 4.43E-10 |
| regulation of transcription by RNA polymerase II | 4.30E-13 | 4.56E-10 |
| positive regulation of nucleic acid-templated transcription | 3.85E-13 | 4.70E-10 |
| regulation of macromolecule biosynthetic process | 5.23E-13 | 5.20E-10 |
| regulation of nucleobase-containing compound metabolic process | 6.35E-13 | 5.61E-10 |
| regulation of RNA metabolic process | 6.21E-13 | 5.81E-10 |
| positive regulation of biological process | 7.03E-13 | 5.88E-10 |
| regulation of nitrogen compound metabolic process | 7.51E-13 | 5.97E-10 |
| regulation of cellular metabolic process | 1.09E-12 | 8.24E-10 |
| positive regulation of RNA metabolic process | 1.25E-12 | 9.02E-10 |
| regulation of cellular macromolecule biosynthetic process | 1.52E-12 | 1.05E-09 |
| regulation of cellular biosynthetic process | 1.59E-12 | 1.06E-09 |
| regulation of primary metabolic process | 2.43E-12 | 1.54E-09 |
