## Supplementary Figure S6 for "Reconstructed signaling and regulatory networks identify potential drugs for SARS-CoV-2 infection"

**A**

Sum threshold

| quantile | 0 | 1 | 2 | 3 | 4 | 5 | 6 |
|---|---|---|---|---|---|---|---|
| 0.01 | 0.521784 | 3.41E-09 | 0.257172 | 1 | 1 | 1 | 1 |
| 0.05 | 0.521784 | 1.12E-09 | 7.47E-05 | 0.087696 | 1 | 1 | 1 |
| 0.1 | 0.521784 | 5.46E-06 | 6.72E-10 | 1.28E-05 | 0.042956 | 0.038893 | 1 |
| 0.15 | 0.521784 | 0.001791 | 1.06E-09 | 5.79E-08 | 0.015509 | 0.13533 | 1 |
| 0.2 | 0.521784 | 0.003871 | 1.11E-10 | 5.77E-09 | 0.00053 | 0.283653 | 1 |
| 0.25 | 0.521784 | 0.012143 | 5.80E-10 | 1.56E-10 | 0.009083 | 0.125431 | 1 |
| 0.3 | 0.521784 | 0.02944 | 6.04E-07 | 1.09E-08 | 0.005948 | 0.29093 | 0.216887 |

**B**

Sum threshold

| quantile | 0 | 1 | 2 | 3 | 4 | 5 | 6 |
|---|---|---|---|---|---|---|---|
| 0.01 | 168 | 37 | 1 | 0 | 0 | 0 | 0 |
| 0.05 | 168 | 96 | 15 | 2 | 0 | 0 | 0 |
| 0.1 | 168 | 113 | 39 | 11 | 2 | 1 | 0 |
| 0.15 | 168 | 114 | 52 | 20 | 4 | 1 | 0 |
| 0.2 | 168 | 125 | 66 | 27 | 8 | 1 | 0 |
| 0.25 | 168 | 128 | 75 | 35 | 8 | 2 | 0 |
| 0.3 | 168 | 130 | 75 | 36 | 10 | 2 | 1 |

**C**

| gene_symbol | hypertension | COPD | diabetes | smoke | cancer | sex | age | sum |
|---|---|---|---|---|---|---|---|---|
| cav1 | -1 | -1 | -1 | -1 | -1 | 0 | 0 | -5 |
| rtn4 | -1 | -1 | -1 | 0 | -1 | 0 | 0 | -4 |
| b4gat1 | 0 | 0 | 0 | -1 | -1 | -1 | -1 | -4 |
| wwp2 | 0 | -1 | -1 | -1 | -1 | 0 | 0 | -4 |
| ube2i | 0 | -1 | -1 | -1 | -1 | 0 | 0 | -4 |
| jun | -1 | 1 | -1 | -1 | -1 | -1 | 0 | -4 |
| stom | -1 | -1 | 1 | -1 | -1 | 0 | 0 | -3 |
| tgfb2 | 0 | 0 | -1 | -1 | -1 | 0 | 0 | -3 |
| erbb2 | 1 | -1 | 0 | -1 | -1 | 0 | -1 | -3 |
| fos | -1 | 0 | 1 | -1 | -1 | 0 | -1 | -3 |
| raly | 0 | 0 | -1 | -1 | 0 | -1 | 0 | -3 |
| acat1 | 0 | -1 | -1 | 0 | 0 | -1 | 0 | -3 |
| hsf1 | -1 | 0 | 0 | -1 | 0 | -1 | 0 | -3 |
| ctnna1 | 0 | -1 | 0 | -1 | -1 | 0 | 0 | -3 |
| brd2 | -1 | 1 | -1 | -1 | -1 | -1 | 1 | -3 |
| nfe2l2 | -1 | -1 | 1 | 0 | -1 | 0 | 0 | -2 |
| myc | -1 | 1 | 0 | 0 | 0 | -1 | -1 | -2 |
| aar2 | -1 | 0 | 0 | -1 | 0 | -1 | 1 | -2 |
| akap8l | 0 | -1 | -1 | 0 | 0 | -1 | 1 | -2 |
| nubp2 | 0 | 0 | -1 | -1 | 0 | 0 | 0 | -2 |
| stx4 | -1 | -1 | 0 | 0 | 0 | 0 | 0 | -2 |
| junb | -1 | 1 | 0 | 0 | -1 | -1 | 0 | -2 |
| snx17 | 0 | -1 | 1 | -1 | -1 | 0 | 0 | -2 |
| jak2 | -1 | 1 | 0 | -1 | -1 | 0 | 0 | -2 |
| ctnna2 | 0 | 0 | -1 | 0 | -1 | 1 | -1 | -2 |
| epha2 | -1 | 1 | -1 | 0 | 0 | -1 | 0 | -2 |
| srf | -1 | 1 | 0 | -1 | -1 | 0 | 0 | -2 |
| dnajb2 | 0 | -1 | 0 | -1 | 0 | 0 | 0 | -2 |
| clu | 1 | -1 | -1 | -1 | -1 | 0 | 1 | -2 |
| plat | -1 | 0 | 0 | -1 | 0 | -1 | 1 | -2 |
| ints1 | 0 | 0 | 1 | 0 | 1 | 0 | 0 | 2 |
| sympk | 0 | 0 | 0 | 0 | 0 | 1 | 1 | 2 |
| ep300 | 0 | 0 | 1 | -1 | 0 | 1 | 1 | 2 |
| il6st | 0 | 1 | 1 | 1 | -1 | 0 | 0 | 2 |
| rab1a | 0 | 1 | 0 | -1 | 1 | 0 | 1 | 2 |
| cul2 | 0 | 1 | 0 | 0 | 0 | 0 | 1 | 2 |
| atm | 1 | 1 | 0 | 1 | -1 | 0 | 0 | 2 |
| atr | 1 | 0 | 0 | 0 | 1 | 0 | 0 | 2 |
| g3bp1 | 0 | 0 | -1 | 1 | 1 | 0 | 1 | 2 |
| cd70 | 0 | 1 | 0 | 0 | 1 | 0 | 0 | 2 |
| os9 | 0 | 0 | 0 | -1 | 1 | 1 | 1 | 2 |
| ints2 | 1 | 1 | 0 | 0 | 0 | 0 | 0 | 2 |
| irak1 | 0 | 0 | 0 | 0 | 1 | 1 | 0 | 2 |
| cs | 0 | 0 | 1 | 0 | 1 | 0 | 0 | 2 |
| lox | -1 | 0 | 1 | 0 | 0 | 1 | 1 | 2 |
| ptprj | -1 | 1 | 1 | 0 | 0 | 1 | 0 | 2 |
| prpf40a | 0 | 0 | 0 | 1 | 1 | -1 | 1 | 2 |
| oat | 1 | 0 | -1 | 0 | 1 | 1 | 0 | 2 |
| mpzl1 | 0 | 1 | -1 | 1 | 1 | 0 | 0 | 2 |
| fgb | -1 | 0 | 1 | 1 | 1 | 1 | -1 | 2 |
| rad18 | 1 | 0 | 0 | 0 | 0 | 0 | 1 | 2 |
| smad3 | 0 | -1 | 1 | -1 | 1 | 1 | 1 | 2 |
| foxm1 | 1 | 0 | 0 | 0 | 1 | 0 | 0 | 2 |
| larp1 | 0 | 0 | 1 | -1 | 1 | 0 | 1 | 2 |
| brca1 | 1 | 0 | 0 | 1 | 1 | 0 | 0 | 3 |
| sox2 | 1 | -1 | 1 | 1 | 1 | 0 | 0 | 3 |
| serpine2 | 1 | 0 | 0 | 0 | 1 | 0 | 1 | 3 |
| zc3h18 | 0 | 1 | 0 | 0 | 0 | 1 | 1 | 3 |
| plau | 0 | 0 | 0 | 1 | 1 | 1 | 0 | 3 |
| ctsb | 1 | 1 | 0 | 1 | -1 | 1 | 0 | 3 |
| itch | 0 | 1 | 0 | 1 | 0 | 0 | 1 | 3 |
| csnk2b | 1 | 1 | 0 | 0 | 1 | 0 | 0 | 3 |
| pfkp | 0 | 1 | 0 | 0 | 1 | 1 | 0 | 3 |
| pdia3 | 0 | 1 | -1 | 0 | 1 | 1 | 1 | 3 |
| nup93 | 0 | 1 | 0 | 0 | 1 | 1 | 1 | 4 |
| neu1 | -1 | 1 | 1 | 1 | 1 | 0 | 1 | 4 |

**D**

| GO biological process complete | raw P value | ▲ FDR |
|---|---|---|
| regulation of protein metabolic process | 1.24E-13 | 1.97E-09 |
| regulation of cellular protein metabolic process | 9.37E-13 | 4.97E-09 |
| response to stress | 1.31E-12 | 5.22E-09 |
| symbiotic process | 2.09E-12 | 5.55E-09 |
| positive regulation of protein metabolic process | 7.24E-13 | 5.75E-09 |
| positive regulation of cellular process | 1.84E-12 | 5.84E-09 |
| response to organic substance | 2.69E-11 | 3.29E-08 |
| regulation of cell population proliferation | 1.72E-11 | 3.42E-08 |
| primary metabolic process | 1.98E-11 | 3.49E-08 |
| regulation of cell communication | 2.68E-11 | 3.56E-08 |
| response to hypoxia | 1.60E-11 | 3.62E-08 |
| viral process | 2.63E-11 | 3.80E-08 |
| response to decreased oxygen levels | 2.56E-11 | 4.06E-08 |
| regulation of signaling | 3.71E-11 | 4.21E-08 |
| response to oxygen levels | 6.02E-11 | 6.38E-08 |
| positive regulation of cellular protein metabolic process | 6.79E-11 | 6.74E-08 |
| positive regulation of biological process | 9.46E-11 | 8.36E-08 |
| regulation of signal transduction | 9.32E-11 | 8.71E-08 |
| positive regulation of cellular metabolic process | 1.26E-10 | 1.06E-07 |
| positive regulation of nitrogen compound metabolic process | 1.37E-10 | 1.09E-07 |
| organic substance metabolic process | 1.46E-10 | 1.11E-07 |
| response to abiotic stimulus | 2.07E-10 | 1.50E-07 |
| positive regulation of metabolic process | 2.20E-10 | 1.52E-07 |
| regulation of protein modification process | 2.42E-10 | 1.60E-07 |
| regulation of cell adhesion | 3.01E-10 | 1.91E-07 |
| regulation of cell motility | 3.41E-10 | 2.09E-07 |
| apoptotic signaling pathway | 3.65E-10 | 2.15E-07 |
| positive regulation of macromolecule metabolic process | 4.87E-10 | 2.77E-07 |
| response to chemical | 5.46E-10 | 2.99E-07 |
