## Supplementary Figure S7 for "Reconstructed signaling and regulatory networks identify potential drugs for SARS-CoV-2 infection"

Top 100 genes + TFs analysis for scRNA-seq SDREM model with protein phosphorylation data integrated

**A**

Sum threshold (columns) × quantile (rows)

| quantile | 0 | 1 | 2 | 3 | 4 | 5 | 6 |
|---|---|---|---|---|---|---|---|
| 0.01 | 0.525038 | 6.33E-06 | 0.201545 | 1 | 1 | 1 | 1 |
| 0.05 | 0.525038 | 1.56E-07 | 1.56E-05 | 0.310931 | 1 | 1 | 1 |
| 0.1 | 0.525038 | 8.51E-07 | 1.03E-10 | 0.000248 | 0.002088 | 0.029587 | 1 |
| 0.15 | 0.525038 | 0.000188 | 1.19E-09 | 2.79E-06 | 0.006142 | 0.104244 | 1 |
| 0.2 | 0.525038 | 0.00477 | 1.36E-09 | 6.93E-07 | 0.012293 | 0.22319 | 1 |
| 0.25 | 0.525038 | 0.007227 | 2.11E-11 | 1.28E-08 | 0.170602 | 0.079134 | 1 |
| 0.3 | 0.525038 | 0.03262 | 1.56E-07 | 8.06E-07 | 0.009743 | 0.195827 | 1 |

**B**

Sum threshold (columns) × quantile (rows)

| quantile | 0 | 1 | 2 | 3 | 4 | 5 | 6 |
|---|---|---|---|---|---|---|---|
| 0.01 | 127 | 25 | 1 | 0 | 0 | 0 | 0 |
| 0.05 | 127 | 72 | 14 | 1 | 0 | 0 | 0 |
| 0.1 | 127 | 96 | 35 | 8 | 3 | 1 | 0 |
| 0.15 | 127 | 98 | 44 | 15 | 4 | 1 | 0 |
| 0.2 | 127 | 98 | 53 | 20 | 5 | 1 | 0 |
| 0.25 | 127 | 103 | 67 | 27 | 4 | 2 | 0 |
| 0.3 | 127 | 101 | 64 | 27 | 8 | 2 | 0 |

**C**

| gene_symbol | hypertension | COPD | diabetes | smoke | cancer | sex | age | sum |
|---|---|---|---|---|---|---|---|---|
| nr3c1 | -1 | -1 | -1 | -1 | -1 | 0 | 0 | -5 |
| cav1 | -1 | -1 | -1 | -1 | -1 | 1 | 0 | -4 |
| jun | -1 | 1 | -1 | -1 | -1 | -1 | 0 | -4 |
| hspa8 | -1 | -1 | 0 | 0 | 0 | -1 | 0 | -3 |
| erbb2 | 1 | -1 | 0 | -1 | -1 | 0 | -1 | -3 |
| srf | -1 | 1 | -1 | -1 | -1 | 0 | 0 | -3 |
| fos | -1 | 0 | 1 | -1 | -1 | 0 | -1 | -3 |
| acat1 | 0 | -1 | -1 | -1 | 0 | -1 | 1 | -3 |
| apc | 0 | -1 | -1 | 0 | -1 | 0 | 0 | -3 |
| mafk | -1 | 0 | 0 | -1 | 0 | 0 | -1 | -3 |
| ctnna1 | 1 | -1 | 0 | -1 | -1 | -1 | 0 | -3 |
| mzf1 | 0 | -1 | 0 | -1 | -1 | 1 | -1 | -3 |
| brd2 | -1 | 1 | -1 | -1 | -1 | -1 | 1 | -3 |
| mcm3ap | 0 | -1 | 0 | -1 | 0 | -1 | 0 | -3 |
| myc | -1 | 1 | 0 | 0 | 0 | -1 | -1 | -2 |
| akap8l | 0 | -1 | -1 | 0 | 0 | -1 | 1 | -2 |
| atf3 | -1 | 1 | 0 | 0 | -1 | 0 | -1 | -2 |
| rxra | 0 | -1 | 0 | -1 | -1 | 0 | 1 | -2 |
| stx4 | -1 | -1 | 0 | 0 | 0 | 0 | 0 | -2 |
| creb1 | 0 | -1 | 1 | 1 | -1 | -1 | -1 | -2 |
| junb | -1 | 1 | 0 | 0 | -1 | -1 | 0 | -2 |
| bach1 | -1 | -1 | 0 | 0 | 0 | 0 | 0 | -2 |
| jak2 | -1 | 1 | 0 | -1 | -1 | 0 | 0 | -2 |
| ccdc8 | 0 | -1 | -1 | 0 | 0 | 0 | 0 | -2 |
| crebbp | 0 | -1 | -1 | -1 | 0 | 0 | 1 | -2 |
| ctnna2 | 0 | 0 | -1 | 0 | -1 | 1 | -1 | -2 |
| irf1 | -1 | 0 | 1 | -1 | -1 | 1 | -1 | -2 |
| stat6 | 0 | 0 | -1 | -1 | -1 | 0 | 1 | -2 |
| rela | -1 | 0 | 0 | 0 | -1 | 1 | -1 | -2 |
| stip1 | -1 | 1 | 1 | 1 | 0 | 0 | 0 | 2 |
| cul2 | 0 | 1 | 0 | 0 | 0 | 0 | 1 | 2 |
| mafg | -1 | 1 | 1 | 0 | 0 | 1 | 0 | 2 |
| atm | 1 | 1 | 0 | 1 | -1 | 0 | 0 | 2 |
| fancg | 0 | 1 | 0 | 0 | 1 | 0 | 0 | 2 |
| mxd1 | -1 | 1 | 0 | 1 | 0 | 0 | 1 | 2 |
| g3bp1 | 0 | 0 | -1 | 1 | 1 | 0 | 1 | 2 |
| esrra | 0 | 1 | -1 | 1 | 1 | 0 | 0 | 2 |
| sumo1 | 1 | 1 | 0 | -1 | 1 | 0 | 0 | 2 |
| cebpg | 0 | 0 | 1 | 1 | 0 | 0 | 0 | 2 |
| fam20c | 0 | 1 | 1 | 0 | 0 | 0 | 0 | 2 |
| hes1 | 1 | 1 | 1 | 1 | -1 | 0 | -1 | 2 |
| elk1 | -1 | -1 | 1 | 1 | 1 | 0 | 1 | 2 |
| irf6 | 1 | -1 | 1 | -1 | 1 | 1 | 0 | 2 |
| arfgef2 | 1 | 0 | -1 | 1 | 1 | -1 | 1 | 2 |
| prpf40a | 0 | 0 | 0 | 1 | 1 | -1 | 1 | 2 |
| hmga1 | -1 | 1 | 0 | 1 | 1 | 0 | 0 | 2 |
| bclaf1 | 0 | -1 | 1 | 1 | 0 | 1 | 0 | 2 |
| tfdp1 | 1 | -1 | 0 | 0 | 1 | 0 | 1 | 2 |
| hnf4a | 0 | 1 | -1 | 1 | 0 | 1 | 0 | 2 |
| smad3 | 0 | -1 | 1 | -1 | 1 | 1 | 1 | 2 |
| dsp | 1 | 1 | -1 | -1 | 1 | 1 | 0 | 2 |
| nr1h3 | 0 | 0 | 1 | 0 | 0 | 0 | 1 | 2 |
| larp1 | 0 | 0 | 1 | -1 | 1 | 0 | 1 | 2 |
| arid5b | 0 | 1 | 0 | 1 | 0 | 0 | 0 | 2 |
| emd | 0 | 1 | 1 | 0 | 1 | 0 | 0 | 3 |
| sympk | 0 | 0 | 0 | 0 | 1 | 1 | 1 | 3 |
| ep300 | 0 | 1 | 1 | -1 | 0 | 1 | 1 | 3 |
| irak1 | 0 | 0 | 1 | -1 | 1 | 1 | 1 | 3 |
| ptprj | -1 | 1 | 1 | 0 | 0 | 1 | 1 | 3 |
| egfr | 0 | 1 | 1 | 1 | 1 | -1 | 0 | 3 |
| neu1 | -1 | 1 | 1 | 1 | 1 | -1 | 1 | 3 |
| fosl2 | -1 | 1 | 0 | 1 | 0 | 1 | 1 | 3 |
| csnk2a2 | 0 | 1 | 1 | -1 | 1 | 0 | 1 | 3 |
| zc3h18 | 0 | 1 | 0 | 0 | 0 | 1 | 1 | 3 |
| itch | 0 | 1 | 0 | 1 | 0 | 0 | 1 | 3 |
| irf5 | 0 | 1 | 0 | 1 | 0 | 0 | 1 | 3 |
| csnk2a1 | 0 | 1 | 1 | 1 | 1 | 0 | 1 | 5 |

**D**

| GO biological process complete | raw P value | FDR |
|---|---|---|
| positive regulation of transcription by RNA polymerase II | 1.51E-21 | 2.39E-17 |
| positive regulation of nitrogen compound metabolic process | 4.95E-21 | 3.93E-17 |
| positive regulation of gene expression | 1.06E-20 | 4.19E-17 |
| positive regulation of macromolecule biosynthetic process | 9.05E-21 | 4.80E-17 |
| positive regulation of cellular biosynthetic process | 5.22E-20 | 1.18E-16 |
| positive regulation of RNA metabolic process | 4.73E-20 | 1.25E-16 |
| positive regulation of biosynthetic process | 9.15E-20 | 1.32E-16 |
| positive regulation of macromolecule metabolic process | 4.24E-20 | 1.35E-16 |
| positive regulation of RNA biosynthetic process | 8.88E-20 | 1.41E-16 |
| positive regulation of nucleic acid-templated transcription | 8.71E-20 | 1.54E-16 |
| regulation of transcription by RNA polymerase II | 8.49E-20 | 1.69E-16 |
| regulation of transcription, DNA-templated | 1.44E-19 | 1.90E-16 |
| positive regulation of transcription, DNA-templated | 1.70E-19 | 1.93E-16 |
| positive regulation of metabolic process | 1.58E-19 | 1.93E-16 |
| regulation of cellular macromolecule biosynthetic process | 3.67E-19 | 3.43E-16 |
| regulation of RNA biosynthetic process | 3.53E-19 | 3.50E-16 |
| regulation of nucleic acid-templated transcription | 3.33E-19 | 3.53E-16 |
| regulation of RNA metabolic process | 8.55E-19 | 7.55E-16 |
| positive regulation of nucleobase-containing compound metabolic process | 1.01E-18 | 8.48E-16 |
| regulation of macromolecule biosynthetic process | 1.41E-18 | 1.12E-15 |
| positive regulation of cellular metabolic process | 3.80E-18 | 2.87E-15 |
| regulation of cellular biosynthetic process | 6.84E-18 | 4.94E-15 |
| regulation of biosynthetic process | 1.54E-17 | 1.02E-14 |
| regulation of nucleobase-containing compound metabolic process | 1.49E-17 | 1.03E-14 |
| positive regulation of cellular process | 2.05E-17 | 1.31E-14 |
| positive regulation of biological process | 3.21E-16 | 1.96E-13 |
| response to stress | 4.61E-16 | 2.61E-13 |
| cellular response to chemical stimulus | 4.60E-16 | 2.71E-13 |
| [illegible] | | |
