## Supplementary Figure S8 for "Reconstructed signaling and regulatory networks identify potential drugs for SARS-CoV-2 infection"

### Top 1000 gene pairs analysis for scRNA-seq SDREM model with protein phosphorylation data integrated

**A**

| | 0 | 1 | 2 | 3 | 4 | 5 | 6 |
|---|---|---|---|---|---|---|---|
| 0.01 | 0.523602 | 1.99E-06 | 0.02703 | 1 | 1 | 1 | 1 |
| 0.05 | 0.523602 | 2.50E-06 | 5.17E-05 | 0.342316 | 1 | 1 | 1 |
| 0.1 | 0.523602 | 2.77E-05 | 4.84E-09 | 0.002397 | 0.032196 | 0.033228 | 1 |
| 0.15 | 0.523602 | 0.001292 | 8.95E-09 | 2.50E-06 | 0.047471 | 0.116503 | 1 |
| 0.2 | 0.523602 | 0.003989 | 2.34E-10 | 1.02E-06 | 0.019243 | 0.247364 | 1 |
| 0.25 | 0.523602 | 0.004596 | 4.50E-11 | 2.67E-09 | 0.012452 | 0.096478 | 1 |
| 0.3 | 0.523602 | 0.016461 | 7.61E-09 | 7.47E-07 | 0.002029 | 0.014349 | 0.018781 |

**B**

| | 0 | 1 | 2 | 3 | 4 | 5 | 6 |
|---|---|---|---|---|---|---|---|
| 0.01 | 143 | 28 | 2 | 0 | 0 | 0 | 0 |
| 0.05 | 143 | 73 | 14 | 1 | 0 | 0 | 0 |
| 0.1 | 143 | 96 | 34 | 7 | 2 | 1 | 0 |
| 0.15 | 143 | 101 | 45 | 16 | 3 | 1 | 0 |
| 0.2 | 143 | 109 | 59 | 21 | 5 | 1 | 0 |
| 0.25 | 143 | 116 | 71 | 30 | 7 | 2 | 0 |
| 0.3 | 143 | 116 | 74 | 29 | 10 | 4 | 2 |

**C**

| gene_symbol | hypertension | COPD | diabetes | smoke | cancer | sex | age | sum |
|---|---|---|---|---|---|---|---|---|
| nr3c1 | -1 | -1 | -1 | -1 | -1 | 0 | 0 | -5 |
| cd81 | 0 | -1 | -1 | -1 | -1 | -1 | 0 | -5 |
| cav1 | -1 | -1 | -1 | -1 | -1 | 1 | 0 | -4 |
| fas | -1 | 0 | -1 | 0 | -1 | -1 | 0 | -4 |
| jun | -1 | 1 | -1 | -1 | -1 | -1 | 0 | -4 |
| hspa8 | -1 | -1 | 0 | 0 | 0 | -1 | 0 | -3 |
| hax1 | -1 | -1 | 0 | -1 | 1 | -1 | 0 | -3 |
| erbb2 | 1 | -1 | 0 | -1 | -1 | 0 | -1 | -3 |
| srf | -1 | 1 | -1 | -1 | -1 | 0 | 0 | -3 |
| fos | -1 | 0 | 1 | -1 | -1 | 0 | -1 | -3 |
| acat1 | 0 | -1 | -1 | -1 | 0 | -1 | 1 | -3 |
| apc | 0 | -1 | -1 | 0 | -1 | 0 | 0 | -3 |
| ctnna1 | 1 | -1 | 0 | -1 | -1 | -1 | 0 | -3 |
| brd2 | -1 | 1 | -1 | -1 | -1 | -1 | 1 | -3 |
| mcm3ap | 0 | -1 | 0 | -1 | 0 | -1 | 0 | -3 |
| scai | 0 | 0 | 0 | 1 | -1 | -1 | -1 | -2 |
| myc | -1 | 1 | 0 | 0 | 0 | -1 | -1 | -2 |
| aar2 | -1 | 0 | 0 | -1 | 0 | -1 | 1 | -2 |
| akap8l | 0 | -1 | -1 | 0 | 0 | -1 | 1 | -2 |
| atf3 | -1 | 1 | 0 | 0 | -1 | 0 | -1 | -2 |
| rxra | 0 | -1 | 0 | -1 | -1 | 0 | 1 | -2 |
| stx4 | -1 | -1 | 0 | 0 | 0 | 0 | 0 | -2 |
| creb1 | 0 | -1 | 1 | 1 | -1 | -1 | -1 | -2 |
| junb | -1 | 1 | 0 | 0 | -1 | -1 | 0 | -2 |
| jak2 | -1 | 1 | 0 | -1 | -1 | 0 | 0 | -2 |
| ccdc8 | 0 | -1 | -1 | 0 | 0 | 0 | 0 | -2 |
| crebbp | 0 | -1 | -1 | -1 | 0 | 0 | 1 | -2 |
| ctnna2 | 0 | 0 | -1 | 0 | -1 | 1 | -1 | -2 |
| rnf213 | -1 | -1 | 0 | 0 | 0 | -1 | 1 | -2 |
| clu | 1 | -1 | -1 | -1 | -1 | 0 | 1 | -2 |
| rela | -1 | 0 | 0 | 0 | -1 | 1 | -1 | -2 |
| stip1 | -1 | 1 | 1 | 1 | 0 | 0 | 0 | 2 |
| cul2 | 0 | 1 | 0 | 0 | 0 | 0 | 1 | 2 |
| atm | 1 | 1 | 0 | 1 | -1 | 0 | 0 | 2 |
| fancg | 0 | 1 | 0 | 0 | 1 | 0 | 0 | 2 |
| sarm1 | 0 | 0 | 0 | 1 | 1 | -1 | 1 | 2 |
| golga5 | 0 | 1 | 1 | 0 | 0 | 0 | 0 | 2 |
| pabpc4 | 0 | 0 | 1 | 0 | 0 | 1 | 0 | 2 |
| g3bp1 | 0 | 0 | -1 | 1 | 1 | 0 | 1 | 2 |
| nr2c2 | 1 | -1 | 0 | 1 | 0 | 0 | 1 | 2 |
| sumo1 | 1 | 1 | 0 | -1 | 1 | 0 | 0 | 2 |
| rif1 | 0 | 1 | 0 | 0 | -1 | 1 | 1 | 2 |
| fam20c | 0 | 1 | 1 | 0 | 0 | 0 | 0 | 2 |
| arfgef2 | 1 | 0 | -1 | 1 | 1 | -1 | 1 | 2 |
| prpf40a | 0 | 0 | 0 | 1 | 1 | -1 | 1 | 2 |
| tut1 | 1 | 1 | 0 | 0 | 0 | 0 | 0 | 2 |
| hmga1 | -1 | 1 | 0 | 1 | 1 | 0 | 0 | 2 |
| ca12 | 1 | 0 | 0 | 1 | 1 | -1 | 0 | 2 |
| bclaf1 | 0 | -1 | 1 | 1 | 0 | 1 | 0 | 2 |
| tfdp1 | 1 | -1 | 0 | 0 | 1 | 0 | 1 | 2 |
| hnf4a | 0 | 1 | -1 | 1 | 0 | 1 | 0 | 2 |
| smad3 | 0 | -1 | 1 | -1 | 1 | 1 | 1 | 2 |
| dsp | 1 | 1 | -1 | -1 | 1 | 1 | 0 | 2 |
| psat1 | 0 | 1 | 1 | 0 | 0 | 0 | 0 | 2 |
| larp1 | 0 | 0 | 1 | -1 | 1 | 0 | 1 | 2 |
| taf4 | 1 | 0 | 0 | 0 | 1 | 0 | 0 | 2 |
| emd | 0 | 1 | 1 | 0 | 1 | 0 | 0 | 3 |
| sympk | 0 | 0 | 0 | 0 | 1 | 1 | 1 | 3 |
| ep300 | 0 | 1 | 1 | -1 | 0 | 1 | 1 | 3 |
| rab1a | 0 | 1 | 1 | -1 | 1 | 0 | 1 | 3 |
| irak1 | 0 | 0 | 1 | -1 | 1 | 1 | 1 | 3 |
| ptprj | -1 | 1 | 1 | 0 | 0 | 1 | 1 | 3 |
| egfr | 0 | 1 | 1 | 1 | 1 | -1 | 0 | 3 |
| neu1 | -1 | 1 | 1 | 1 | 1 | -1 | 1 | 3 |
| csnk2a2 | 0 | 1 | 1 | -1 | 1 | 0 | 1 | 3 |
| zc3h18 | 0 | 1 | 0 | 0 | 0 | 1 | 1 | 3 |
| fgb | -1 | 1 | 1 | 1 | 1 | 1 | -1 | 3 |
| plau | 0 | 0 | 0 | 1 | 1 | 1 | 0 | 3 |
| itch | 0 | 1 | 0 | 1 | 0 | 0 | 1 | 3 |
| ssbp1 | 0 | 1 | 1 | 1 | 1 | 0 | 0 | 4 |
| bet1 | 1 | 1 | 0 | 0 | 1 | 0 | 1 | 4 |

**D**

| GO biological process complete | raw P value | FDR |
|---|---|---|
| response to stress | 4.38E-14 | 6.96E-10 |
| symbiotic process | 1.53E-13 | 8.12E-10 |
| positive regulation of nitrogen compound metabolic process | 1.14E-13 | 9.09E-10 |
| positive regulation of cellular process | 2.39E-13 | 9.50E-10 |
| positive regulation of metabolic process | 3.48E-13 | 1.11E-09 |
| positive regulation of cellular metabolic process | 7.15E-13 | 1.62E-09 |
| positive regulation of macromolecule metabolic process | 6.42E-13 | 1.70E-09 |
| viral process | 1.64E-12 | 3.27E-09 |
| response to organic substance | 3.24E-12 | 5.72E-09 |
| positive regulation of biological process | 3.99E-12 | 6.34E-09 |
| regulation of cell communication | 4.92E-12 | 7.12E-09 |
| regulation of signaling | 6.97E-12 | 9.24E-09 |
| positive regulation of gene expression | 8.10E-12 | 9.90E-09 |
| regulation of signal transduction | 1.25E-11 | 1.32E-08 |
| organic substance metabolic process | 1.18E-11 | 1.34E-08 |
| macromolecule metabolic process | 2.48E-11 | 2.46E-08 |
| regulation of response to stimulus | 2.96E-11 | 2.62E-08 |
| gene expression | 2.83E-11 | 2.65E-08 |
| cellular response to chemical stimulus | 3.46E-11 | 2.90E-08 |
| positive regulation of transcription by RNA polymerase II | 4.96E-11 | 3.94E-08 |
| gland development | 5.33E-11 | 4.03E-08 |
| cellular response to stress | 6.53E-11 | 4.72E-08 |
| cellular response to organic substance | 8.04E-11 | 5.56E-08 |
| primary metabolic process | 1.08E-10 | 6.63E-08 |
| cellular metabolic process | 1.08E-10 | 6.89E-08 |
| metabolic process | 1.07E-10 | 7.07E-08 |
| interspecies interaction between organisms | 1.37E-10 | 8.07E-08 |
| response to chemical | 1.63E-10 | 9.25E-08 |
| regulation of apoptotic process | 1.99E-10 | 1.05E-07 |
| positive regulation of cellular biosynthetic process | 1.93E-10 | 1.06E-07 |
| regulation of cell death | 2.45E-10 | 1.25E-07 |
