## Supplementary Figure S9 for "Reconstructed signaling and regulatory networks identify potential drugs for SARS-CoV-2 infection"

**A**

Sum threshold

| quantile | 0 | 1 | 2 | 3 | 4 | 5 | 6 |
|---|---|---|---|---|---|---|---|
| 0.01 | 0.525237 | 0.000581 | 0.198729 | 1 | 1 | 1 | 1 |
| 0.05 | 0.525237 | 5.35E-08 | 3.05E-06 | 1 | 1 | 1 | 1 |
| 0.1 | 0.525237 | 1.11E-05 | 2.97E-09 | 1.21E-07 | 0.02523 | 0.029131 | 1 |
| 0.15 | 0.525237 | 0.00463 | 5.84E-09 | 2.33E-06 | 0.034246 | 0.1027 | 1 |
| 0.2 | 0.525237 | 0.014111 | 8.74E-10 | 1.42E-07 | 0.011567 | 1 | 1 |
| 0.25 | 0.525237 | 0.006658 | 1.67E-09 | 3.46E-08 | 0.006407 | 0.365551 | 1 |
| 0.3 | 0.525237 | 0.043158 | 3.67E-07 | 2.04E-08 | 4.98E-05 | 0.047722 | 1 |

**B**

Sum threshold

| | 0 | 1 | 2 | 3 | 4 | 5 | 6 |
|---|---|---|---|---|---|---|---|
| 0.01 | 125 | 20 | 1 | 0 | 0 | 0 | 0 |
| 0.05 | 125 | 73 | 15 | 0 | 0 | 0 | 0 |
| 0.1 | 125 | 89 | 32 | 12 | 2 | 1 | 0 |
| 0.15 | 125 | 86 | 42 | 15 | 3 | 1 | 0 |
| 0.2 | 125 | 92 | 53 | 21 | 5 | 0 | 0 |
| 0.25 | 125 | 102 | 61 | 26 | 7 | 1 | 0 |
| 0.3 | 125 | 98 | 62 | 30 | 12 | 3 | 0 |

**C**

| gene_symbol | hypertension | COPD | diabetes | smoke | cancer | sex | age | sum |
|---|---|---|---|---|---|---|---|---|
| cebpd | -1 | 0 | -1 | 0 | -1 | -1 | 0 | -4 |
| foxo3 | -1 | -1 | -1 | -1 | -1 | 0 | 1 | -4 |
| jun | -1 | 1 | -1 | -1 | -1 | -1 | 0 | -4 |
| egr1 | -1 | 0 | 0 | -1 | -1 | 0 | 0 | -3 |
| nr3c1 | 0 | 0 | -1 | -1 | -1 | 0 | 0 | -3 |
| erbb2 | 1 | -1 | 0 | -1 | -1 | 0 | -1 | -3 |
| fos | -1 | 0 | 1 | -1 | -1 | 0 | -1 | -3 |
| raly | 0 | 0 | -1 | -1 | 0 | -1 | 0 | -3 |
| acat1 | 0 | -1 | -1 | 0 | 0 | -1 | 0 | -3 |
| ctnna1 | 0 | -1 | 0 | -1 | -1 | 0 | 0 | -3 |
| nfe2l2 | -1 | -1 | 1 | 0 | -1 | 0 | 0 | -2 |
| myc | -1 | 1 | 0 | 0 | 0 | -1 | -1 | -2 |
| jund | -1 | 0 | 0 | 0 | -1 | 0 | 0 | -2 |
| rora | -1 | -1 | 0 | 0 | 0 | 0 | 0 | -2 |
| pbx1 | 1 | -1 | 0 | -1 | -1 | 0 | 0 | -2 |
| akap8l | 0 | -1 | -1 | 0 | 0 | -1 | 1 | -2 |
| atf3 | -1 | 1 | 0 | 0 | -1 | 0 | -1 | -2 |
| rxra | 0 | -1 | 0 | -1 | -1 | 0 | 1 | -2 |
| stx4 | -1 | -1 | 0 | 0 | 0 | 0 | 0 | -2 |
| junb | -1 | 1 | 0 | 0 | -1 | -1 | 0 | -2 |
| meis1 | 0 | -1 | 0 | -1 | -1 | 0 | 1 | -2 |
| mef2a | 0 | 0 | 0 | -1 | -1 | 0 | 0 | -2 |
| stat6 | 0 | 0 | -1 | -1 | -1 | 0 | 1 | -2 |
| rpl36 | 0 | 0 | 0 | -1 | 0 | 0 | -1 | -2 |
| clu | 1 | -1 | -1 | -1 | -1 | 0 | 1 | -2 |
| sympk | 0 | 0 | 0 | 0 | 0 | 1 | 1 | 2 |
| ep300 | 0 | 0 | 1 | -1 | 0 | 1 | 1 | 2 |
| cul2 | 0 | 1 | 0 | 0 | 0 | 0 | 1 | 2 |
| atm | 1 | 1 | 0 | 1 | -1 | 0 | 0 | 2 |
| atr | 1 | 0 | 0 | 0 | 1 | 0 | 0 | 2 |
| elk4 | 0 | 0 | 1 | 1 | 0 | 1 | -1 | 2 |
| g3bp1 | 0 | 0 | -1 | 1 | 1 | 0 | 1 | 2 |
| ckap4 | 0 | 0 | -1 | 0 | 1 | 1 | 1 | 2 |
| rif1 | 0 | 1 | 0 | 0 | -1 | 1 | 1 | 2 |
| dek | 0 | 0 | 0 | 0 | 0 | 1 | 1 | 2 |
| atf7 | 1 | 1 | 0 | 1 | 0 | 0 | -1 | 2 |
| ptprj | -1 | 1 | 1 | 0 | 0 | 1 | 0 | 2 |
| prpf40a | 0 | 0 | 0 | 1 | 1 | -1 | 1 | 2 |
| tfap2a | 1 | -1 | 0 | 1 | 1 | -1 | 1 | 2 |
| oat | 1 | 0 | -1 | 0 | 1 | 1 | 0 | 2 |
| smad3 | 0 | -1 | 1 | -1 | 1 | 1 | 1 | 2 |
| larp1 | 0 | 0 | 1 | -1 | 1 | 0 | 1 | 2 |
| hif1a | 0 | 1 | -1 | 1 | 1 | 1 | 0 | 3 |
| brca1 | 1 | 0 | 0 | 1 | 1 | 0 | 0 | 3 |
| zc3h18 | 0 | 1 | 0 | 0 | 0 | 1 | 1 | 3 |
| plau | 0 | 0 | 0 | 1 | 1 | 1 | 0 | 3 |
| dsp | 1 | 1 | 0 | -1 | 1 | 1 | 0 | 3 |
| itch | 0 | 1 | 0 | 1 | 0 | 0 | 1 | 3 |
| yy1 | 0 | -1 | 0 | 1 | 1 | 1 | 1 | 3 |
| csnk2b | 1 | 1 | 0 | 0 | 1 | 0 | 0 | 3 |
| pdia3 | 0 | 1 | -1 | 0 | 1 | 1 | 1 | 3 |
| tmpo | 1 | -1 | 1 | 1 | 1 | 1 | 0 | 4 |
| sp1 | 1 | 1 | 1 | 0 | 0 | 1 | 0 | 4 |

**D**

| GO biological process complete | raw P value | FDR |
|---|---|---|
| positive regulation of macromolecule metabolic process | 2.53E-19 | 4.00E-15 |
| positive regulation of metabolic process | 5.52E-18 | 2.91E-14 |
| positive regulation of nitrogen compound metabolic process | 3.73E-18 | 2.95E-14 |
| positive regulation of transcription by RNA polymerase II | 9.99E-18 | 3.95E-14 |
| positive regulation of cellular biosynthetic process | 1.98E-17 | 6.26E-14 |
| positive regulation of biosynthetic process | 3.29E-17 | 7.43E-14 |
| positive regulation of cellular metabolic process | 2.98E-17 | 7.85E-14 |
| positive regulation of macromolecule biosynthetic process | 4.38E-17 | 8.66E-14 |
| positive regulation of nucleobase-containing compound metabolic process | 1.38E-16 | 2.42E-13 |
| positive regulation of RNA biosynthetic process | 1.39E-15 | 1.69E-12 |
| positive regulation of transcription, DNA-templated | 1.37E-15 | 1.80E-12 |
| positive regulation of RNA metabolic process | 1.21E-15 | 1.91E-12 |
| positive regulation of nucleic acid-templated transcription | 1.37E-15 | 1.97E-12 |
| regulation of macromolecule metabolic process | 5.78E-15 | 6.52E-12 |
| regulation of transcription by RNA polymerase II | 6.22E-15 | 6.56E-12 |
| regulation of cellular macromolecule biosynthetic process | 1.47E-14 | 1.45E-11 |
| regulation of macromolecule biosynthetic process | 1.92E-14 | 1.78E-11 |
| regulation of RNA metabolic process | 2.26E-14 | 1.98E-11 |
| regulation of nucleobase-containing compound metabolic process | 2.92E-14 | 2.43E-11 |
| regulation of nitrogen compound metabolic process | 5.37E-14 | 4.25E-11 |
