## Supplementary Figure S10 for "Reconstructed signaling and regulatory networks identify potential drugs for SARS-CoV-2 infection"

**A**

Sum threshold

| quantile | 0 | 1 | 2 | 3 | 4 | 5 | 6 |
|---|---|---|---|---|---|---|---|
| 0.01 | 0.521593 | 1.52E-08 | 0.03733 | 1 | 1 | 1 | 1 |
| 0.05 | 0.521593 | 3.70E-10 | 7.06E-06 | 0.393861 | 1 | 1 | 1 |
| 0.1 | 0.521593 | 8.16E-05 | 3.12E-12 | 2.68E-06 | 0.044327 | 0.03957 | 1 |
| 0.15 | 0.521593 | 0.000992 | 6.14E-12 | 7.27E-10 | 0.003046 | 0.137563 | 1 |
| 0.2 | 0.521593 | 0.001939 | 2.09E-13 | 8.04E-14 | 3.56E-06 | 0.28789 | 1 |
| 0.25 | 0.521593 | 0.004611 | 2.73E-12 | 6.66E-11 | 1.30E-05 | 0.025304 | 1 |
| 0.3 | 0.521593 | 0.010853 | 2.67E-09 | 1.90E-10 | 6.14E-05 | 0.005354 | 0.220283 |

**B**

Sum threshold

| | 0 | 1 | 2 | 3 | 4 | 5 | 6 |
|---|---|---|---|---|---|---|---|
| 0.01 | 171 | 36 | 2 | 0 | 0 | 0 | 0 |
| 0.05 | 171 | 99 | 17 | 1 | 0 | 0 | 0 |
| 0.1 | 171 | 107 | 44 | 12 | 2 | 1 | 0 |
| 0.15 | 171 | 118 | 58 | 23 | 5 | 1 | 0 |
| 0.2 | 171 | 130 | 74 | 35 | 11 | 1 | 0 |
| 0.25 | 171 | 135 | 83 | 36 | 13 | 3 | 0 |
| 0.3 | 171 | 138 | 85 | 40 | 14 | 5 | 1 |

**C**

| gene_symbol | hypertension | COPD | diabetes | smoke | cancer | sex | age | sum |
|---|---|---|---|---|---|---|---|---|
| cav1 | -1 | -1 | -1 | -1 | -1 | 0 | 0 | -5 |
| rtn4 | -1 | -1 | -1 | 0 | -1 | 0 | 0 | -4 |
| b4gat1 | 0 | 0 | 0 | -1 | -1 | -1 | -1 | -4 |
| foxo3 | -1 | -1 | -1 | -1 | -1 | 0 | 1 | -4 |
| wwp2 | 0 | -1 | -1 | -1 | -1 | 0 | 0 | -4 |
| cd81 | 0 | -1 | -1 | -1 | -1 | 0 | 0 | -4 |
| jun | -1 | 1 | -1 | -1 | -1 | -1 | 0 | -4 |
| nr3c1 | 0 | 0 | -1 | -1 | -1 | 0 | 0 | -3 |
| fkbp8 | -1 | -1 | -1 | 1 | 0 | -1 | 0 | -3 |
| erbb2 | 1 | -1 | 0 | -1 | -1 | 0 | -1 | -3 |
| fos | -1 | 0 | 1 | -1 | -1 | 0 | -1 | -3 |
| raly | 0 | 0 | -1 | -1 | 0 | -1 | 0 | -3 |
| scap | 0 | -1 | 0 | -1 | -1 | 0 | 0 | -3 |
| ncaph2 | -1 | -1 | -1 | -1 | 0 | 1 | 0 | -3 |
| ctnna1 | 0 | -1 | 0 | -1 | -1 | 0 | 0 | -3 |
| fanci | 1 | 1 | 0 | 0 | 1 | -1 | 1 | 3 |
| hif1a | 0 | 1 | -1 | 1 | 1 | 1 | 0 | 3 |
| brca1 | 1 | 0 | 0 | 1 | 1 | 0 | 0 | 3 |
| bet1 | 0 | 1 | 0 | 0 | 1 | 0 | 1 | 3 |
| atp1b1 | 1 | 0 | 1 | 1 | 1 | 0 | -1 | 3 |
| serpine2 | 1 | 0 | 0 | 0 | 1 | 0 | 1 | 3 |
| ca12 | 1 | 0 | 0 | 1 | 1 | 0 | 0 | 3 |
| zc3h18 | 0 | 1 | 0 | 0 | 0 | 1 | 1 | 3 |
| plau | 0 | 0 | 0 | 1 | 1 | 1 | 0 | 3 |
| dsp | 1 | 1 | 0 | -1 | 1 | 1 | 0 | 3 |
| ctsb | 1 | 1 | 0 | 1 | -1 | 1 | 0 | 3 |
| itch | 0 | 1 | 0 | 1 | 0 | 0 | 1 | 3 |
| nup188 | 0 | 1 | 0 | 1 | 0 | 0 | 1 | 3 |
| yy1 | 0 | -1 | 0 | 1 | 1 | 1 | 1 | 3 |
| csnk2b | 1 | 1 | 0 | 0 | 1 | 0 | 0 | 3 |
| pdia3 | 0 | 1 | -1 | 0 | 1 | 1 | 1 | 3 |
| nup93 | 0 | 1 | 0 | 0 | 1 | 1 | 1 | 4 |
| neu1 | -1 | 1 | 1 | 1 | 1 | 0 | 1 | 4 |
| tmpo | 1 | -1 | 1 | 1 | 1 | 1 | 0 | 4 |
| sp1 | 1 | 1 | 1 | 0 | 0 | 1 | 0 | 4 |

**D**

| GO biological process complete | raw P value | FDR |
|---|---|---|
| regulation of multicellular organismal process | 1.47E-09 | 1.16E-05 |
| regulation of pri-miRNA transcription by RNA polymerase II | 1.42E-09 | 2.24E-05 |
| response to hormone | 2.41E-08 | 7.63E-05 |
| positive regulation of pri-miRNA transcription by RNA polymerase II | 2.39E-08 | 9.46E-05 |
| response to endogenous stimulus | 2.16E-08 | 1.14E-04 |
| regulation of multicellular organismal development | 7.59E-08 | 2.00E-04 |
| response to abiotic stimulus | 1.43E-07 | 3.22E-04 |
| regulation of cellular component movement | 2.90E-07 | 5.10E-04 |
| viral life cycle | 2.71E-07 | 5.36E-04 |
| biological process involved in symbiotic interaction | 9.39E-07 | 1.48E-03 |
| regulation of cell motility | 1.11E-06 | 1.59E-03 |
| viral entry into host cell | 2.05E-06 | 2.16E-03 |
| response to stress | 1.81E-06 | 2.20E-03 |
| regulation of locomotion | 1.70E-06 | 2.24E-03 |
| response to ketone | 2.28E-06 | 2.25E-03 |
| negative regulation of multicellular organismal process | 2.00E-06 | 2.26E-03 |
| entry into host | 2.89E-06 | 2.68E-03 |
| response to hypoxia | 3.95E-06 | 3.47E-03 |
