## Supplementary Table S8 for "Reconstructed signaling and regulatory networks identify potential drugs for SARS-CoV-2 infection": readme.txt

These tables show the gene differential expression using different quantile level threshod, which is shown at the end each table name. For example, 0.01 means that 
genes with quantile > 0.99 are highly expressed with label of 1, genes with quantile < 0.01 are lowly expressed with label of -1, and
all the other genes are non differentially expressed with label of 0.
